# GT61 β-1,2-xylosyltransferases define a conserved xylan modification in gymnosperm and Arabidopsis primary cell walls

**DOI:** 10.1101/2025.08.06.668652

**Authors:** Henry Temple, Yoshihisa Yoshimi, Katharine Davis, Theodora Tryfona, Aleksandra Liszka, Henry Yates, James Andrew London, Alberto Echevarría-Poza, Joel Wurman-Rodrick, Li Yu, Glenn Thorlby, Nikolaj Spodsberg, Kyrin R Hanning, Christina Fleischmann, Xiaolan Yu, Katherine Stott, Kristian B. R. M. Krogh, Mathias Soreiul, Nadine Anders, Paul Dupree

## Abstract

Plant primary and secondary cell walls differ in molecular composition, structure, and mechanical properties. While secondary wall xylan has been extensively characterised, the structure of xylan in primary walls remains less well understood, particularly in gymnosperms. Here, we identify a previously uncharacterised β-1,2-linked xylosyl side chain in conifer and *Arabidopsis thaliana* xylan. Using enzymatic fingerprinting, NMR, and mass spectrometry, we show that this structure is positioned two xylose residues away from glucuronic acid substitutions, forming an evenly patterned substituted xylan. This spacing pattern is consistent with xylan–cellulose interaction, suggesting a structural role in primary wall architecture.

This modification, found in primary wall-rich tissues of diverse conifer species, including needles and pro-embryogenic mass (PEM), is also present in Arabidopsis callus. We demonstrate that conifer Group III GT61 glycosyltransferases introduce this modification with consistent positional specificity. In Arabidopsis, three closely related GT61 enzymes act redundantly to generate the same structure, and their combined loss results in its complete absence. These findings uncover a conserved primary wall xylan modification in seed plants and define the GT61 enzymes responsible for its biosynthesis, opening new avenues to explore how xylan structure contributes to primary wall function.

**Significance Statement:** Xylan structure is well characterised in secondary walls, but its primary wall counterpart remains poorly understood. We identified a conserved β-1,2-xylosyl modification on xylan in the primary walls of conifers and Arabidopsis. This side chain is positioned at a defined position from a glucuronic acid substitution and is introduced by GT61 glycosyltransferases that cluster in one phylogenetic subclade. Our findings revealed a previously unrecognised xylan structural pattern and the biosynthetic enzymes responsible for its addition. This work expands the current understanding of primary wall architecture across seed plants.

## Introduction

Xylan is a polymer of β-(1→4)-linked D-xylose that forms a major component of plant cell walls. Its conformation and interactions with other matrix polymers are shaped by the identity and distribution of its side-chain substitutions (Busse-Wicher, Grantham, *et al*., 2016; Terrett and Dupree, 2019). These modifications vary across species, tissues, and developmental stages (Curry *et al*., 2023), influencing xylan’s folding, accessibility, and capacity to bind to cellulose or other wall components.

In secondary cell walls (SCWs), xylan substitution patterns are relatively well characterised. Both grasses and most gymnosperms produce glucuronoarabinoxylan (GAX), in which xylosyl residues can carry α-1,3-L-arabinofuranose (Araf) and α-1,2-D-[4-O-methyl] glucuronic acid ([Me]GlcA) side chains, albeit with distinct patterns in each group (Tryfona *et al*., 2023; Busse-Wicher, Li, *et al*., 2016). In the three main groups of gymnosperms (Conifer, Gingko, and Cycad), wood contains xylan that lacks acetylation, but is modified by 3-linked arabinosyl and 2-linked methylated glucuronyl side chains in an even-substitution pattern, which has been shown to be essential for xylan–cellulose interactions (Busse-Wicher, Li, *et al*., 2016; Martínez-Abad *et al*., 2017; Grantham *et al*., 2017). In contrast, eudicot SCWs contain acetylated glucuronoxylan (GX), which entirely lacks arabinosylation (Rennie and Scheller, 2014; Scheller and Ulvskov, 2010).

The substitution patterns of xylan in primary cell walls (PCWs) remain poorly resolved, in part because PCW xylan is typically low in abundance and often masked by the dominant SCW, a limitation that is particularly pronounced in gymnosperms, where, to our knowledge, no primary wall xylan structures have yet been reported. Nonetheless, it is known that PCWs of Arabidopsis young stems and callus contain xylan with arabinopyranosyl-modified, GlcA-containing oligosaccharides (XU^Arap,[Me]^XXX) (Mortimer *et al*., 2015; Yu *et al*., 2024). In addition, young, non-lignified grass tissues have been reported to accumulate more densely substituted GAX than their lignified counterparts. Moreover, based on linkage analysis, 2-linked xylosyl residues have been proposed in Arabidopsis seed mucilage (Voiniciuc *et al*., 2015; Ralet *et al*., 2016). While this points to additional branching in primary wall xylan, the precise nature and positioning of these residues within the polymer cannot be fully determined from linkage data alone. These observations suggest that xylan in primary walls may harbour distinct substitution patterns with potential functional implications, yet detailed structural characterisation remains limited, particularly in gymnosperms, where no primary wall xylan structures have previously been described.

Pentosyl substitutions on the xylan backbone are catalysed by glycosyltransferases from the GT61 family. The first characterised members of this family, a wheat and two rice α-1-3-arabinofuranosyltransferases (XATs), introduce Ara*f* side chains onto xylan in grasses (Anders *et al*., 2012). Since then, additional GT61 biosynthetic activities have been identified, including β-1,2-xylosyltransferases (XYXTs) from rice and *Pinus taeda* (Zhong *et al*., 2018; Zhong *et al*., 2022), but the existence and tissue location of any corresponding β-1,2-xylosylated structures have yet to be clearly described in rice and conifers.

Here, we characterised a previously undescribed β-1,2-linked xylosyl side chain on glucuronoxylan in the primary cell walls of both conifers and Arabidopsis. Using enzymatic fingerprinting, Matrix-Assisted Laser Desorption/Ionization Collision-Induced Dissociation (MALDI-CID) mass spectrometry, and Nuclear Magnetic Resonance (NMR) spectroscopy, we identified an even-substitution patterned xylan containing a characteristic oligosaccharide in which β-1,2-xylosyl branches are consistently positioned two xylose residues apart from GlcA substitutions. This modification is synthesised by a conserved sub-clade of GT61 glycosyltransferases: Group III members in conifers and the Arabidopsis homologues AtXYXT1 (also known as MUCI21/MUM5), AtXYXT2, and AtXYXT3. These enzymes exhibit positional specificity, introducing xylosyl branches in a defined spatial relationship to GlcA. Our findings define a previously unrecognised structural motif in plant primary walls, identify its biosynthetic origin, and demonstrate that β-1,2-xylosylation is a conserved feature of primary cell wall xylan at least in conifers and Arabidopsis.

## Results

### Conifer xylan can be modified by 2-linked xylosyl side chains that are sensitive to arabinofuranosidase *Pa*GH62

To analyse the xylan in conifer tissues with different proportions of PCW and SCWs, we isolated alcohol insoluble material (AIR) from needles, which contains both wall types (Michavila *et al*., 2025), and compared it to AIR derived from wood, which consists predominantly of SCW. We selected three closely related *Pinaceae* species as cell wall sources, *Pinus radiata*, *Picea abies* and *Pseudotsuga menziesii* (Douglas fir), and one species of the more distantly related species, *Metasequoia glyptostroboides* (Metasequoia). AIR was hydrolysed with *Neocallimastix patriciarum* GH11 xylanase (GH11), and the resulting xylo-oligosaccharides were analysed using PolyAcrylamide Carbohydrate gel Electrophoresis (PACE). As expected, all samples yielded common products such as xylose, xylobiose, and GlcA-modified oligosaccharides XUXX and XUUXX. However, the banding pattern differed subtly between wood (W) and needles (N), particularly among higher molecular weight, substituted oligosaccharides migrating between markers X_5_-X_6_ (Supporting Figure 1A). In particular the relative proportion between the xylo-oligosaccharide A, running just below the marker of X_6_, and xylo-oligosaccharide, running just above X_5_, is shifted towards Y being relatively more abundant in needles and A being relatively more abundant in wood.

To characterise the structure of the two xylo-oligosaccharides (A and Y) further, we analysed *Picea abies* and Metasequoia AIR extracted from needles with GH11 and other enzymes that hydrolyse xylan side chains: *Bacteroides ovatus* α-glucuronidase GH115 (GH115), *Meripilus giganteus* α-arabinofuranosidase GH51 (GH51) and *Penicillium aurantiogriseum* α-arabinofuranosidase GH62 (GH62). GH115 removes GlcA side chains from the xylan backbone, and when used in combination with GH11, both bands A and Y are sensitive to digestion, resulting in the formation of xylo-oligosaccharides A′ and Y′, which lack GlcA substitutions (Figure 1A, B). Additional hydrolysis with the arabinofuranosidase GH51 and GH62, showed that A is sensitive to both GH51 and GH62, confirming Ara*f* side chains on the xylo-oligosaccharide A. This is consistent with published structures of xylan in SCW, producing a xylo-oligosaccharide substituted by Ara*f* and GlcA and has been previously characterised as: XA^3^XU^2^XX (Cresswell *et al*., 2025). In contrast, the band Y from needles is resistant to GH51 hydrolysis, but not to GH62 hydrolysis (Figure 1A, B). Both enzymes GH51 and GH62 have been described to hydrolyse 2- and 3-linked arabinofuranosyl side chains of the xylan backbone, although GH51 is not specific for xylan (Beylot *et al*., 2001; Wilkens *et al*., 2017). To understand this unexpected difference in enzyme activity of GH51 and GH62 arabinofuranosidases, we analysed the detailed structure of the GH51-resistant Y band. We isolated the GH11 and GH51 hydrolysis product (Y oligosaccharide) from *Metasequioa* needles using size-exclusion chromatography and pooled fractions in which the xylo-oligosaccharide was enriched. After 2-aminobenzoic acid (2-AA) labelling, we subjected the oligosaccharide to mass spectrometry and MALDI-CID analysis. The detected mass of 1276 *m/z* of the xylo-oligosaccharide is consistent with MeGlcAPentose_7_-2AA. Further analysis of the fragmentation ions of the xylo-oligosaccharide reveals a backbone of 6 xyloses, substituted at the −3 position with a pentose (P) and at the −5 position with a methylated GlcA side chain (U): XU^4m2^XPXX (Figure 1C). Although the mass is consistent with the oligosaccharide XA^3^XU^2^XX (band A), the order of substitutions is not, which likely accounts for their distinct migration in the gel. However, the oligosaccharide is resistant to GH51 hydrolysis, which may point to a different nature of the pentose side chain, and which remained unresolved by MALDI-CID analysis.

**Figure 1:**
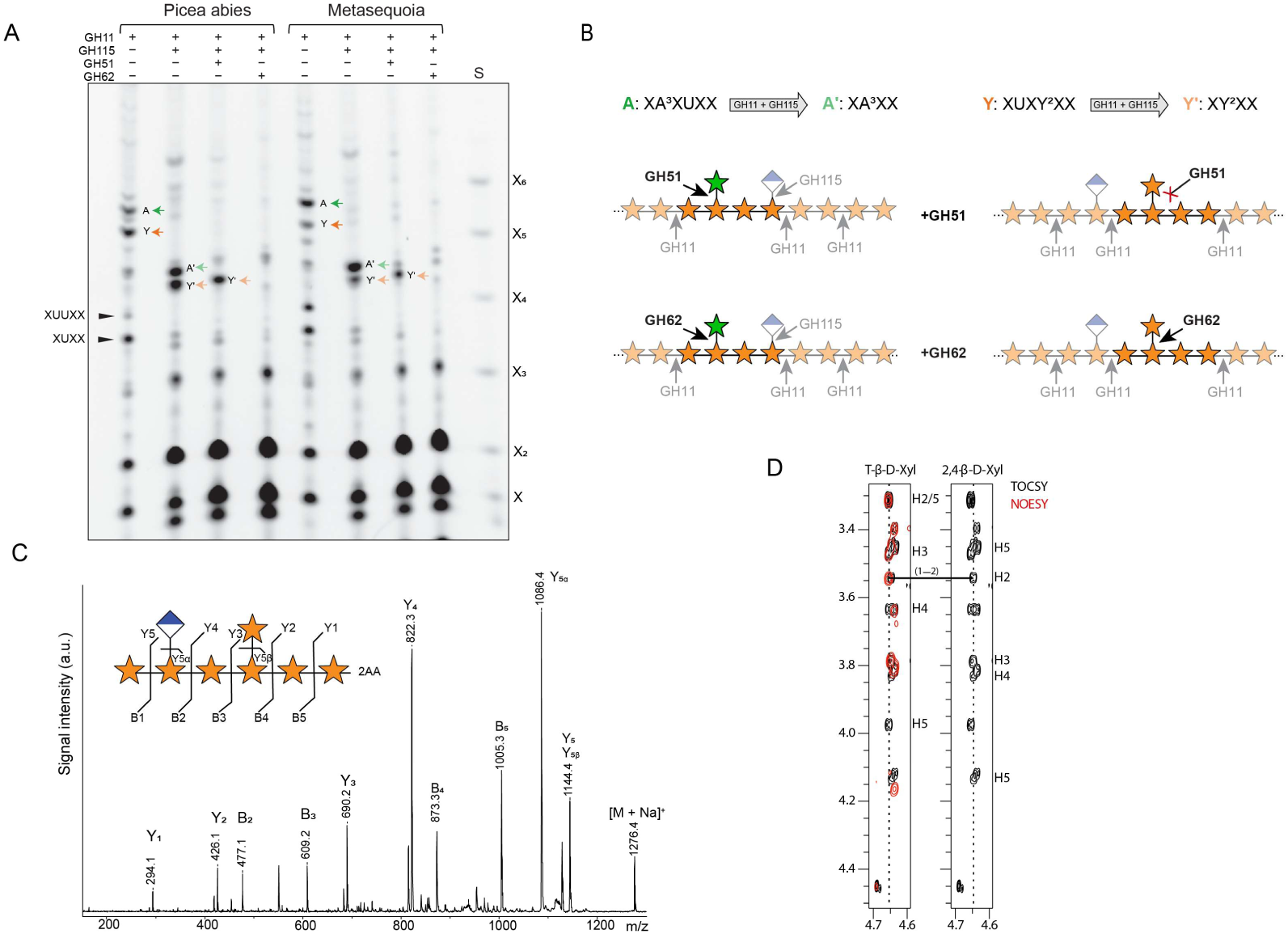
Novel 1,2-xylosylated xylo-oligosaccharide in conifer needles. **A** PACE analysis of AIR from needles of *Picea abies* and *Metasequoia glyptostroboides*. AIR was hydrolysed with xylanase GH11, glucuronidase GH115 and arabinofuranosidase GH51 or GH62 as indicated. S: Standard is the xylose ladder X-X_6_. A: XUXA^3^XX, Y: XUXY^2^XX, A’: XA^3^XX, Y’: XY^2^XX. **B** Graphical representation of hydrolysis reactions shown in PACE gel in figure 1A. AIR hydrolysed by GH11 alone produces oligosaccharide A and Y, which is hydrolysed by the combination of GH11 and GH115 to A’ and Y’ (top). Depending on the arabinofuranosidase (GH51 or GH62) used and the pentosyl side chain present the glycosidic bond of the side chain is hydrolysed (black arrow) or maintained (red cross). Note GH11 and GH115 hydrolysis reactions are greyed out. Symbol Nomenclature for Glycans used (Varki *et al*., 2015). **C** MALDI CID analysis of 2-AA-labelled xylo-oligosaccharide from needles of *Metasequoia* glyptostroboides, showing a pentosyl substitution (here shown as xylose) at −3 and a glucuronyl substitution at −5. **D** Solution NMR analysis of the unlabelled xylo-oligosaccharide from needles of *Metasequoia* glyptostroboides, showing the T-β-D-Xyl-(1 → 2)-β-D-Xyl side chain to backbone linkage, using *1*H strip plots for *1*H,*1*H-TOCSY (*black*) and *1*H,*1*H-NOESY (*red*) spectra to highlight the NOE connectivity arising from this glycosidic linkage, confirming the oligosaccharide Y as XUXY^2^XX.

We used solution-state NMR to further characterise the enriched xylo-oligosaccharide’s structure. 2D *1*H,*13*C-heteronuclear single-quantum coherence (HSQC) spectrum assignment, using two-dimensional experiments, indicated a series of xylose, arabinose, and 4-O-methylated glucuronic acid residues were present within the samples tested. Comparative analyses of 2D *1*H,*1*H-total correlation spectroscopy (TOCSY) and *1*H,*1*H-nuclear overhauer effect spectroscopy (NOESY) spectra, to assess the linkages present, highlighted several xyloses that exhibited linkages between their anomeric and position-4 hydroxyl groups, consistent with a hexameric backbone chain (Supporting Figure S2). Further, linkages between a MeGlcA residue’s and a xylose residue’s anomeric hydroxyl groups and the position-2 hydroxyl groups of the −5 and −3 backbone xyloses, respectively, indicated both were present as branching side chains (Supporting Figure S2 and Figure 1D). No linkages could be detected between the arabinose residues present or from these residues to either xylose or 4-O-methyl glucuronic acid residues. The assignments made and linkage analyses performed indicated a XU^4m2^XY^2^XX (Y^2^ referring to β-1,2-linked xylosyl side chains) structure was present in needles. This configuration explains the oligosaccharide’s resistance to GH51, which hydrolyses arabinofuranosyl side chains. Surprisingly, however, it was sensitive to GH62, indicating that *Pa*GH62, under our conditions, can also cleave β-1,2-linked xylosyl side chains, whereas GH51 remains specific to arabinose.

Together, these NMR and enzymatic analyses confirm that the oligosaccharide carries a β-1,2-linked xylosyl side chain and reveal a defined, even-patterned modification on conifer xylan that is enriched in needles compared to wood.

### Xylosyl substitutions in conifer primary cell walls occur at a defined distance from GlcA substitutions, consistent with the enzymatic specificity of group III GT61s

Having identified the β-1,2-xylosylated oligosaccharide XUXY^2^XX enriched in conifer needles, we next asked whether this structure is broadly associated with primary wall xylan across tissues. We generated a collection of *Pinus radiata* tissues including pro-embryogenic masses (PEM), needles, and herbaceous stem. These tissues were then analysed for GT61 gene expression and corresponding xylan substitution patterns. Cell wall materials were hydrolysed using GH11 xylanase, with or without GH115, GH51, and GH62. As expected, XA^3^XUXX (oligosaccharide A) was abundant in herbaceous stem and showed sensitivity to GH51 and GH62, consistent with arabinofuranosyl side chains. In needles, XA^3^XUXX was less prominent, and completely absent in PEM. Instead, the GH51-resistant XUXY^2^XX (oligosaccharide Y) dominated (Figure 2A), suggesting that this structure is a characteristic of gymnosperm primary wall xylan.

**Figure 2:**
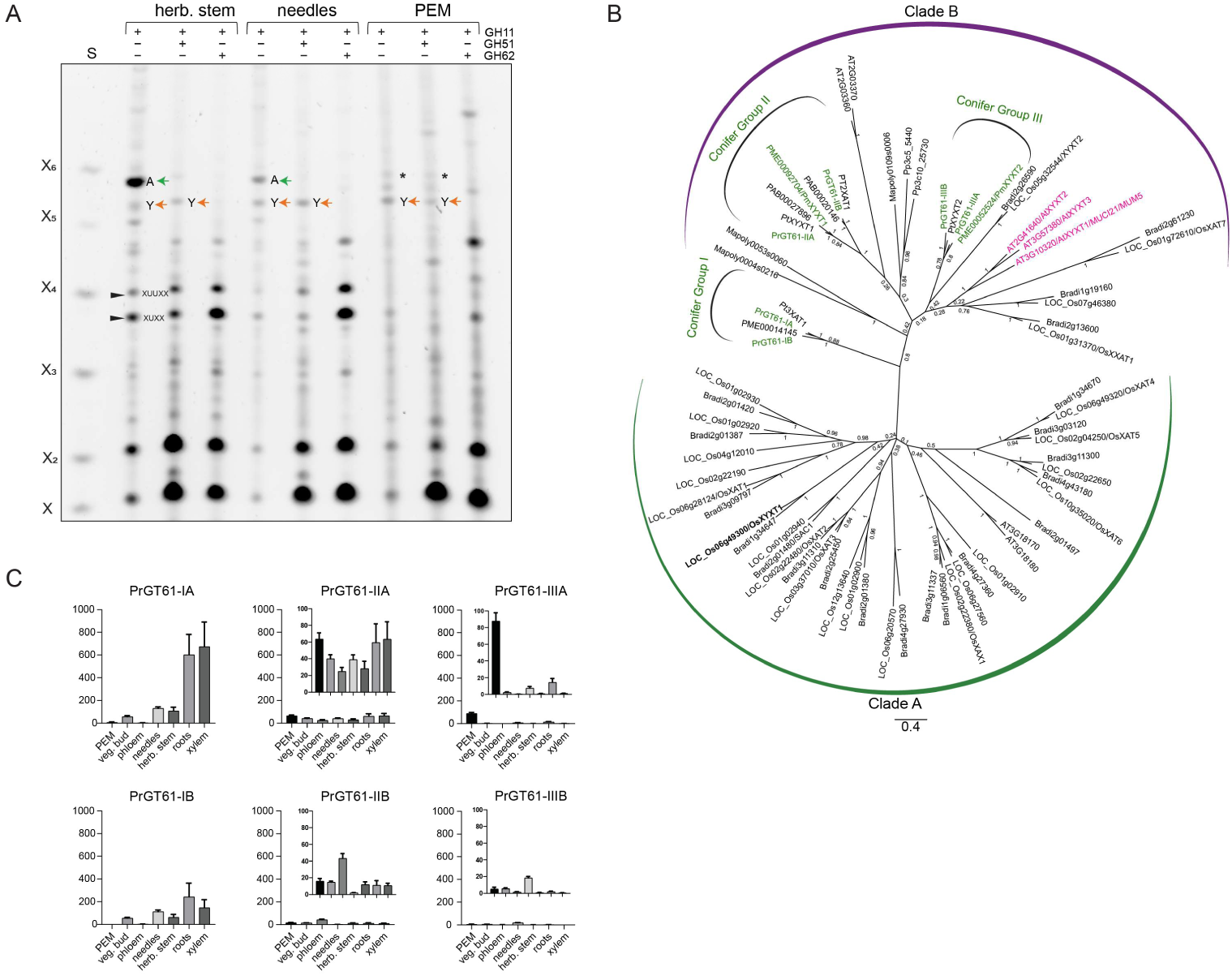
Consistent with gene expression xylan β-1,2 xylosytransferase is present in primary cell wall-rich tissues. **A** PACE analysis of AIR from herbaceous stem (herb. stem), needles and pro-embryogenic masses (PEM) from *Pinus radiata*. AIR was hydrolysed with xylanase GH11, arabinofuranosidase GH51 or GH62 as indicated. Note the change of relative abundance of the oligosaccharides XA^3^XUXX (A) and XUXY^2^XX (Y) in the different tissues. S: Standard is the xylose ladder X-X_6_. Asterisks (*) represents background band, insensitive to GH62. **B** Phylogenetic analysis of full-length GT61 proteins, showing the three conifer groups I, II and III. Douglas fir PmXYXT1 and PmXYXT2 are highlighted in green, Arabidopsis AtXYXT1, AtXYXT2 and AtXYXT3 are highlighted in magenta. OsXYXT1 highlighted in bold. **C** Absolute quantification of expression of six *Pinus radiata* GT61s from conifer group I (PrGT61-IA and PrGT61-IB), group II (PrGT61-IIA and PrGT61-IIB) and group III (PrGT61-IIIA and PrGT61-IIIB) using droplet digital ddPCR in different tissue. Tissues shown are: PEM, vegetative bud (veg. bud), phloem, needles, roots, herbaceaus stem (herb stem) and xylem. Lowly expressed genes are shown with different scale on Y-axis as inserts. Error bars represent the standard deviation from mean. Note the specific expression of PrGT61-IIIA in PEMs.

To identify the enzymes responsible for the β-1,2-xylosylation detected in primary wall xylan, we studied the GT61 family, which includes glycosyltransferases involved in arabinose and xylose substitution of the xylan backbone (Anders *et al*., 2012; Zhong *et al*., 2018). Xylan β-1,2-xylosyltransferases (XYXTs) are members of this family, characterised by their ability to transfer xylosyl residues to the xylan backbone producing a β-1,2-linkages. Zhong *et al*., 2022 proposed three major GT61 conifer groups based on phylogeny and function, with group I comprising α-1,3-arabinosyltransferases (3-XATs), group II including both β-1,2-xylosyltransferase (XYXT) and α-1,2-arabinosyltransferase (2-XAT) activities, and group III containing only XYXTs. This framework provided a useful basis for investigating which GT61 enzymes may be responsible for the specific β-1,2-xylosylation pattern we observed in gymnosperm primary cell walls.

We constructed a phylogeny incorporating GT61 sequences from multiple conifer species including six unreported *P. radiata* genes, which grouped consistently with the classification established by Zhong *et al*., 2022 (Figure 2B). This, combined with our matched tissue collection and xylan oligosaccharide profiling, enabled us to associate GT61 gene expression patterns with specific xylan cell wall types (Figure 2A–C). Group I genes (PrGT61-IA and PrGT61-IB) were strongly expressed in herbaceous stem and xylem which are tissues enriched in secondary walls, while very low abundant in PEM which are PCW exclusive. This pattern is consistent with the enrichment of XA^3^XUXX in wood and absence in PEM, suggesting 3-XATs are involved in secondary wall xylan biosynthesis. Group II genes (PrGT61-IIA and PrGT61-IIB) were expressed more broadly across tissues, including needles and PEM but lacked strong tissue-specific enrichment. In contrast, group III gene PrGT61-IIIA showed a specifically strong expression in PEM, coinciding with the accumulation of the GH51-resistant, GH62-sensitive oligosaccharide XUXY^2^XX. Based on this correlation and its phylogenetic placement, we prioritised *Pr*GT61-IIIA as a candidate XYXT responsible for β-1,2-xylosylation in primary wall xylan.

To investigate the biosynthetic activity of conifer GT61s, we turned to *Pseudotsuga menziesii*, which, in our analysis, contains only one full-length GT61 gene per group, allowing straightforward selection for functional characterisation. We cloned and characterised PmXYXT2 (group III) and PmXYXT1 (group II) as potential XYXTs. These are the respective orthologues of *P. taeda* PtXYXT2 and PtXYXT1, which were independently shown by Zhong et al. (2022), during the course of this work, to encode β-1,2-xylosyltransferases.

We began by assessing the activity of PmXYXT2, as a candidate representative of group III. PmXYXT2-GFP was transiently expressed in *Nicotiana benthamiana*, where it showed Golgi localisation, consistent with a role in hemicellulose biosynthesis (Supplementary Figure S4). Microsome-enriched fractions were incubated with UDP-xylose and deacetylated Arabidopsis wild-type xylan as the acceptor substrate (Figure 3A, Supporting Figure S3A). Digestion of the reaction products with GH11 xylanase yielded an oligosaccharide that co-migrated with XUXY^2^XX (oligosaccharide Y). Given the use of UDP-xylose as donor, and the product’s resistance to GH51 but sensitivity to GH62, the incorporated sugar is most likely a β-1,2-linked xylosyl side chain, consistent with the enzymatic digestion patterns previously observed for this structure. When the reaction products were digested with GH11 and GH115 together, a band migrating as XY^2^XX (Y′ oligosaccharide) appeared, consistent with removal of GlcA allowing GH11 access to the backbone. The absence of Y′ in GH11-only digests suggests that PmXYXT2 adds a xylosyl side chain near GlcA, which initially blocks GH11 access. This positional specificity mirrors the defined spacing seen in native XUXY^2^XX-GH11 product from gymnosperm primary wall–rich tissues.

**Figure 3:**
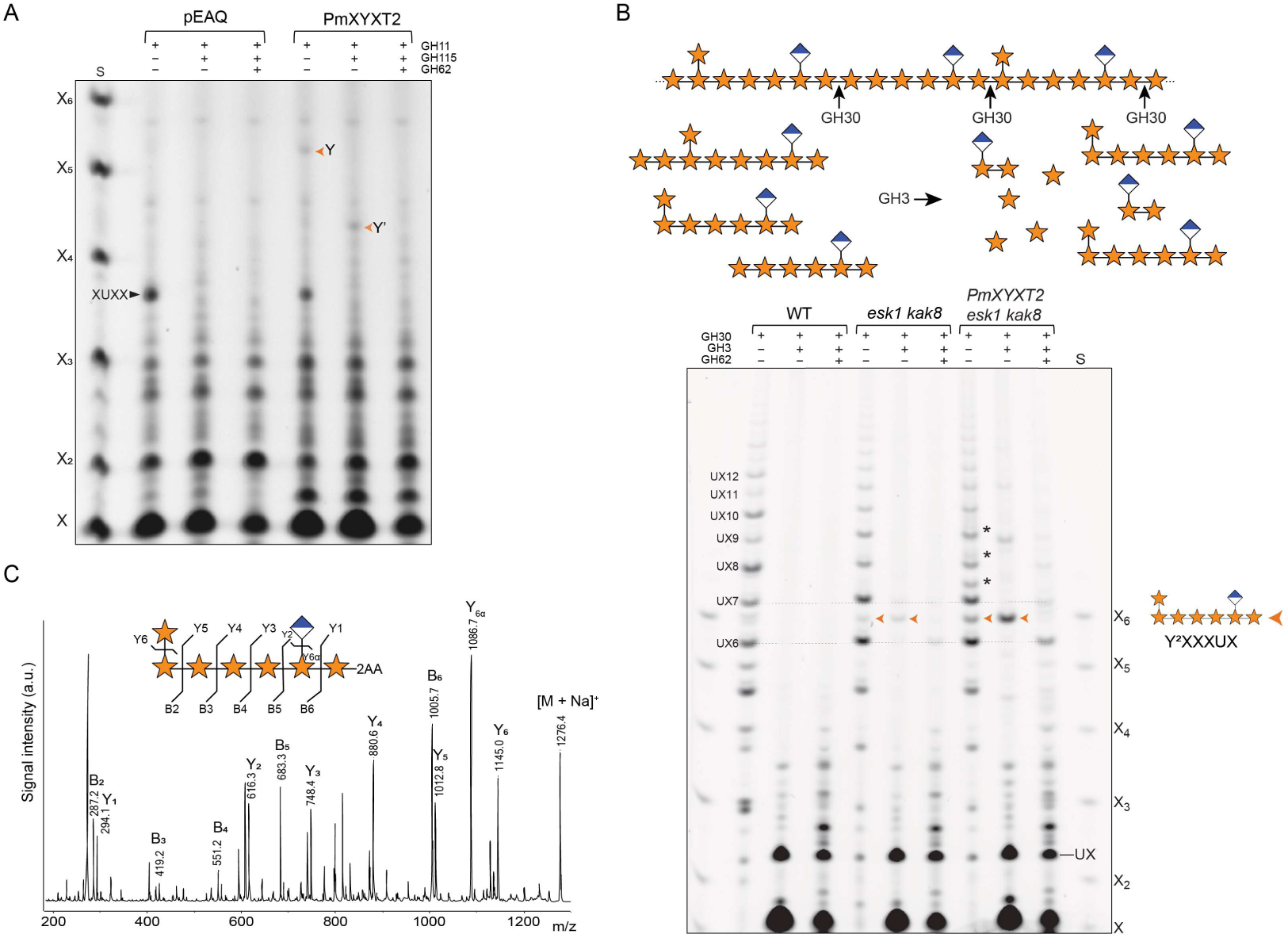
*In vitro*- and ectopic activity of *Pseudotsuga menziesii* xylan β-1,2 xylosytransferase PmXYXT2 shows even patterning with GlcA side chains. **A** *In vitro* activity of tobacco microsomes expressing PmXYXT2 using de-acetylated Arabidopsis wildtype xylan as acceptor. Reaction products were hydrolysed by xylanase GH11, glucuronidase GH115 and GH62 as indicated and analysed by PACE. S: Standard is the xylose ladder X-X_6_. Y: XUXY^2^XX, Y’: XY^2^XX. pEAQ correspond to empty vector. **B** Graphical representation of the hydrolysis reactions by GH30 of xylan modified by PmXYT2 in Arabidopsis *esk1 kak8* (top). GH30 cleaves at the +2 position relative to GlcA substitutions, while GH3 removes terminal xylose residues from the non-reducing end until blocked by a backbone substitution. PACE analysis of PmXYXT2 activity ectopically expressed in *esk1 kak8* (bottom). AIR from basal stems of wild-type (WT), *esk1 kak8*, and *esk1 kak8* expressing *PmXYXT2* under the *IRX3* promoter was hydrolysed using GH30 alone, GH30 in combination with GH3, or GH30 with GH3 followed—after enzyme removal—by GH62. S: Xylose ladder standard (X–X6). Y^2^XXXUX is indicated by an orange arrow. Black asterisks indicate UXn oligosaccharides carrying additional substitutions from PmXYXT2. The collapse of these into a single major oligosaccharide after GH3 treatment suggests a repeating unit that is halted by a backbone substitution. **C** MALDI CID analysis of 2-AA-labelled major xylo-oligosaccharide produced by PmXYXT2 expressed in *esk1 kak8*, hydrolysed with GH30 and GH3, showing the positions of side chains characterised by PACE as Y^2^XXXUX.

In contrast, after GH11 digestion alone, PmXYXT1 (group II) (Supporting Figure S3A) produced both XUXY^2^XX and XY^2^XX, the latter being a structure not detected in our PACE analysis of conifer primary wall-rich tissues. This suggests that PmXYXT1-mediated xylosylation is not strictly restricted to a defined distance from GlcA. Upon GH115 treatment, the intensity of the XY^2^XX band increased, revealing that additional xylosylated fragments became accessible to xylanase GH11 once GlcA was removed. These findings suggest that PmXYXT1 can place β-1,2-linked xylosyl side chains near GlcA residues, but without a consistent or defined spatial relationship. Compared to PmXYXT2, which adds xylosyl branches at a preferred distance from GlcA, PmXYXT1 shows a more flexible and heterogeneous substitution pattern.

To validate these findings *in planta*, we heterologously expressed both enzymes under the secondary wall–specific *AtIRX3* promoter in *Arabidopsis thaliana* WT, *gux1 gux2* (which lacks GlcA, (Mortimer *et al*., 2010)), and *esk1 kak8* (reduced acetylation, (Bensussan *et al*., 2015)). PmXYXT2 was active in the *esk1 kak8* background but only marginally active in WT and *gux1 gux2* (Supporting Figure S3B). This indicates that PmXYXT2 is more active under low-acetylation conditions, consistent with the unacetylated nature of conifer xylan (Supporting Figure S3B). In contrast, PmXYXT1 was most active in *gux1 gux2*, weakly active in *esk1 kak8*, and inactive in WT (Supporting Figure S3C), suggesting that GlcA has a stronger effect in preventing PmXYXT1 activity on xylan than acetylation.

To further investigate the relationship between PmXYXT2 activity and the positioning of GlcA substitutions, we hydrolysed SCW AIR from Arabidopsis *esk1 kak8* plants expressing PmXYXT2 using *Erwinia chrysanthemi* GH30 glucuronoxylanase (GH30), which is a GlcA appendage-dependant xylanase cleaving at the +2 position relative to GlcA (Hurlbert and Preston, 2001; St. John *et al*., 2006; Bromley *et al*., 2013; Yu *et al*., 2024). Subsequently we used *Trichoderma reesei* GH3 β-xylosidase (GH3), which removes xyloses from the non-reducing end until blocked by substitution (Figure 3B and C) (Tenkanen *et al*., 1996). In WT, GH30 digestion produced the expected ladder of mainly even-patterned-glucuronidated oligosaccharides ranging from UX_5_ to at least UX_16_, as described in Bromley *et al*., 2013. In *esk1 kak8*, the same oligosaccharides were observed, but the dominance of the even patterning of GlcA substitutions was attenuated, as previously reported (Grantham *et al*., 2017).

Upon GH3 digestion, the UX_n_ ladder collapsed in both WT and *esk1 kak8*, yielding primarily xylose and UX. However, in *esk1 kak8*, a faint new band remained between UX_6_ and UX_7,_ which was absent in WT and was sensitive to GH62 hydrolysis. After GH62 hydrolysis the oligosaccharide co-migrates with UX_6_, indicating that the band between UX_6_ and UX_7_ corresponds to UX_6_ containing a GH62-sensitive-pentose modification. When PmXYXT2 was expressed in the *esk1 kak8* background, new oligos appeared in between the UX_n_ ladder. Following GH3 digestion, most of these bands collapsed, with a prominent increase in the same GH62-sensitive band observed in *esk1 kak8* (Figure 3B). Based on migration and enzymatic sensitivity, the main band was interpreted as a UX_6_ oligosaccharide carrying one GlcA and a single pentose substitution.

To confirm its structure, the band was analysed by MALDI-CID MS. The resulting fragmentation confirmed a structure with one GlcA and a single pentose side chain at the non-reducing end of the molecule, consistent with YXXXUX (Figure 3C). GH3 digestion cleaved the xylan backbone completely up to the terminal xylose carrying the pentose branch, consistent with a 2-linked substitution. If the pentose had been 3-linked, GH3 would likely have stalled one residue earlier, as substitutions at position 3 sterically hinder the enzyme from accessing the preceding glycosidic bond (Tenkanen *et al*., 1996). Together, these results confirm the β-1,2-linkage and support a repeating backbone structure of even-patterned substitutions such as Y^2^XXXUXY^2^XXXUX, where the underlined segment corresponds to the oligosaccharide released by GH11 digestion observed in PCW-enriched conifer tissues.

Together with the xylanase GH11-derived signature of XUXY^2^XX in gymnosperm primary wall-rich tissues, these findings provide strong support that PmXYXT2 is the β-1,2-xylosyltransferase responsible for the evenly spaced GlcA–Xyl motif in *Pseudotsuga* primary cell wall xylan.

### The specific β-1,2-xylosylation of primary cell wall xylan is conserved in Arabidopsis and mediated by GT61 enzymes

The presence of a faint oligosaccharide co-migrating with Y^2^XXXUX in cell wall material digested with GH30 and GH3 of *esk1 kak8* Arabidopsis stems (orange arrow heads) (Figure 3B) suggested that β-1,2-xylosyltransferase activity may also exist in eudicots. We hypothesised that, as in conifers, this activity may be greater in primary cell walls. To investigate this, we analysed AIR from Arabidopsis stem, leaf, and callus tissues following GH11 digestion in combination with GH51 and GH62 (Figure 4A). While no GH62-sensitive bands were detected in WT stems, leaf material showed a very faint GH62-sensitive, and GH51-resistant band migrating at a similar position as XUXY^2^XX from conifers. This band was substantially stronger in callus, which consists of primary walls, supporting the idea that this structure is more frequent in PCW xylan.

**Figure 4:**
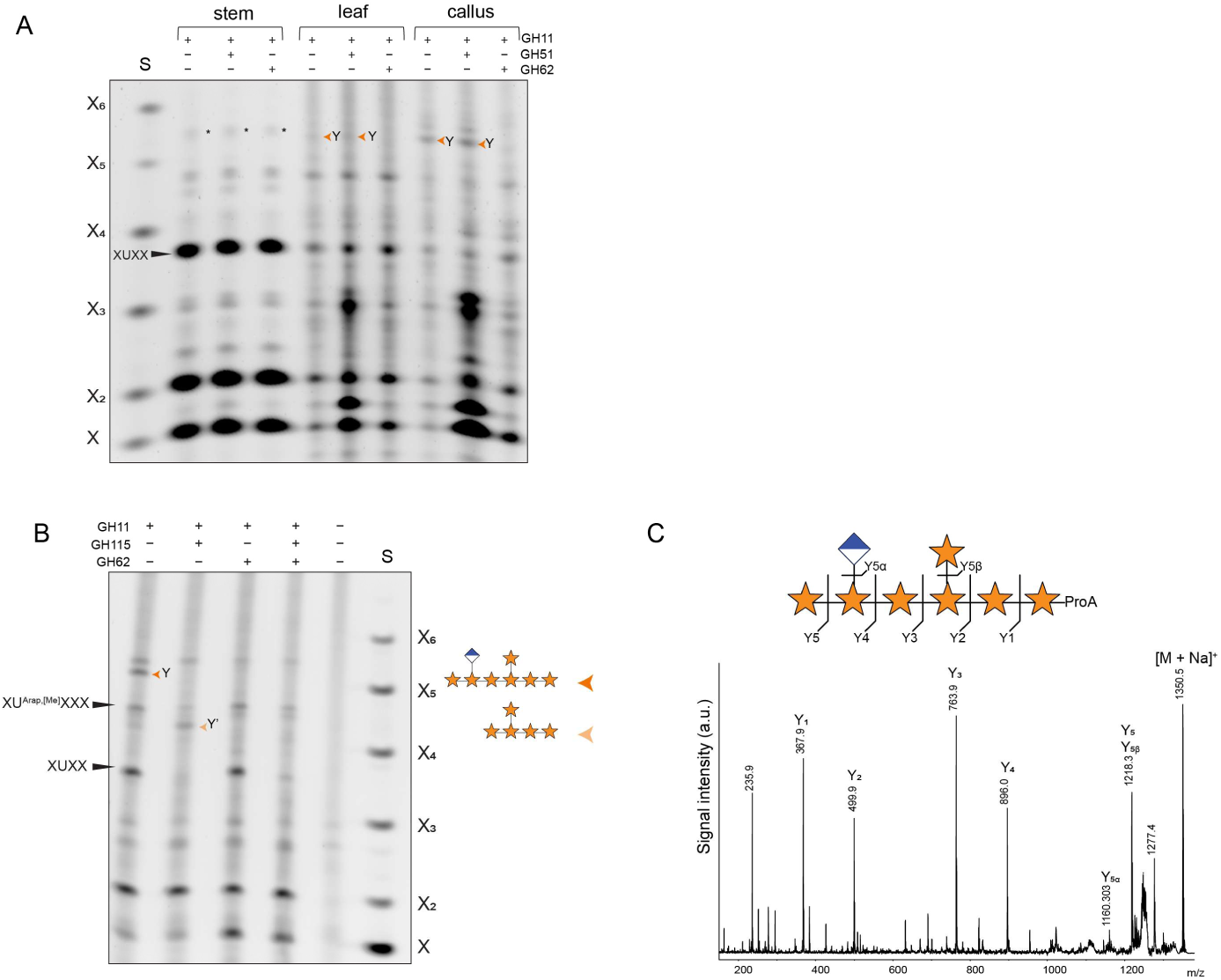
Xylosyl side chains are conserved in Arabidopsis primary cell walls. **A** PACE analysis of *Arabidopsis thaliana* stem, leaf and callus derived AIR hydrolysed with GH11, GH51 and GH62 as indicated. Note XUXY^2^XX (Y) is mostly detectable in callus. Asterisks mark background bands not sensitive to GH62 hydrolysis. S: Standard is the xylose ladder X-X_6_. **B** PACE analysis of *Arabidopsis thaliana* AIR derived from callus hydrolysed with GH11, GH115 and GH62 as indicated. **C** MALDI CID analysis of enriched ProA-laballed xylo-oligosaccharide from Arabidopsis callus generated by GH11 hydrolysis, showing the positions of side chains characterised by PACE as XUXYXX.

To further characterise this GH62-sensitive band, we performed additional xylan fingerprinting in Arabidopsis callus (Figure 4B). Digestion with GH11 and GH115 glucuronidase shifted the band to a position consistent with XY^2^XX, following removal of GlcA. Treatment with GH11 and GH62 hydrolysed the XUXY^2^XX to XUXX. Triple digestion with GH11, GH115, and GH62 eliminated both the GH62-sensitive band and XUXX, leaving only the previously described XU^Ara*p*,[Me]^XXX oligosaccharide (Mortimer *et al*., 2015) which is resistant to GH115 hydrolysis (Yu *et al*., 2024).

To confirm the structure, the digest was labelled with procainamide (ProA) and analysed by MALDI-CID mass spectrometry (Figure 4C). The detected mass of 1350 m/z is consistent with a MeGlcAPentose7–ProA composition and the resulting fragmentation profile verified the oligosaccharide as XUXPXX. The oligosaccharide’s resistance to GH51 and GH62 sensitivity, combined with its migration pattern matching the conifer XUXY^2^XX, we conclude that both gymnosperm and Arabidopsis primary cell wall xylans exhibit a conserved β-1,2-xylosylation pattern. This modification occurs at defined positions relative to GlcA, producing an evenly spaced substitution, compatible with xylan–cellulose interactions.

To identify the responsible enzymes for β-1,2-linked xylose branches in Arabidopsis, we looked at the homologous genes to conifer group III. Phylogenetic analysis places MUCI21/MUM5 (At3g10320), along with its close homologues At2g41640 and At3g57380, within GT61 clade B—as closest Arabidopsis homologs to the conifer group III β-1,2-xylosyltransferases. Although MUCI21/MUM5 has previously been implicated in xylan branching based on glycosidic linkage analysis (Voiniciuc *et al*., 2015; Ralet *et al*., 2016), direct enzymatic evidence for β-1,2-xylosyltransferase activity in Arabidopsis has been lacking.

To test this, we expressed MUCI21/MUM5 under the SCW *AtIRX3* promoter in the *esk1 kak8* background and compared its activity to PmXYXT2 and the previously characterised OsXYXT1 (Zhong *et al*., 2018). Hydrolysis of Arabidopsis basal stem AIR with GH11 and GH115 glucuronidase generated oligosaccharides migrating as XY^2^XX (Y) in all three transgenic lines,consistent with xylan carrying β-1,2-linked xylosyl side chains (Figure 5A). This is confirmed by the band being resistant to GH51 hydrolysis, but fully digestable by GH62, diagnostic for β-1,2-linked xylosyl substitution (Figure 5A, Supporting Figure S5A). GH11 hydrolysis of Metasequoia wood xylan yielded mainly XA^3^XX, which was fully sensitive to GH51, confirming arabinose substitution (Figure 5A). Interestingly, OsXYXT1 produced a more complex oligosaccharide pattern, indicating densely substituted xylan (Figure 5A, Supporting Figure S5A). This contrasts with the limited set of GH62-sensitive oligosaccharides generated by MUCI21/MUM5 and PmXyXT2, suggesting that these two xylosyltransferases have stricter positional specificity. OsXYXT1 branching behaviour is reminiscent of PmXYXT1 (Supporting Figure S3), suggesting mechanistic differences among β-1,2-xylosyltransferases. Based on its ability to introduce GH62-sensitive, GH51-resistant β-1,2-linked xylosyl residues onto xylan— comparable to other experimentally validated XYXTs—we conclude that MUCI21/MUM5 encodes a β-1,2-xylosyltransferase. We therefore designate it *Arabidopsis thaliana* xylan xylosyltransferase 1 (AtXYXT1), based on its activity and phylogenetic relationship to other XYXTs.

**Figure 5:**
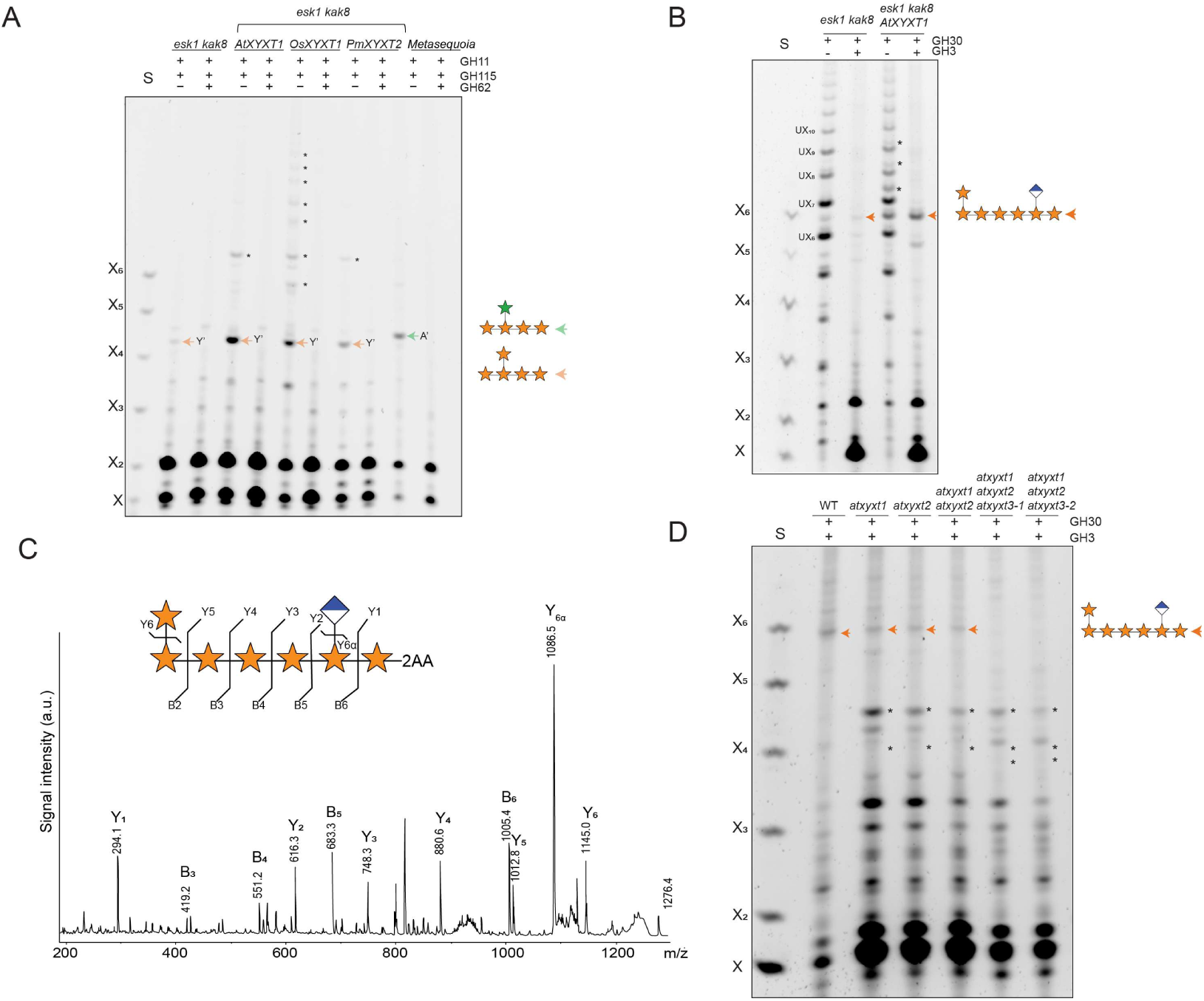
AtXYXT1 acts as β-1,2-xylosyltransferases in Arabidopsis primary cell walls. **A** PACE analysis of *esk1 kak8* xylan expressing ATXYXT1/MUCI21/MUM5, OsXYXT1 and PmXYXT2 under the *IRX3* promoter. AIR from stem was hydrolysed by GH11, GH115 and GH62 as indicated. Control: *Metasequoia* glyptostroboides wood. Note, that XY^2^XX and XA^3^XX run differently in a PACE gel. Asterisks indicate additional, unidentified oligosaccharides that likely represent more complex xylosylated xylan structures. S: Standard is the xylose ladder X-X_6_. **B** AIR of stem of *esk1 kak8* expressing AtXYXT1 under the *IRX3* promoter was hydrolysed by GH11 and GH3 Y^2^XXXUX (orange arrow). Black asterisks indicate UXn oligosaccharides carrying additional substitutions from AtXYXT1. **C** MALDI-CID analysis of 2-AA-labelled xylo-oligosaccharide Y^2^XXXUX produced by AtXYXT1 expressed in *esk1 kak8*, hydrolysed with GH30 and GH3. Note, this is the same oligosaccharide produced by PmXYXT2. **D** PACE analysis of *atxyxt* mutants. GH30 and GH3 hydrolysis of AIR callus from Arabidopsis *atxyxt1* and atxyxt2 single mutants, *atxyxt1* atxyxt2 double mutant, and two triple mutants *atxyxt1* atxyxt2 *atxyxt3-1 and atxyxt1* atxyxt2 *atxyxt3-2*. Oligosaccharide Y^2^XXXUX = orange arrow. Reduction of the Y^2^XXXUX band in the single and double mutants, and its complete absence in the triple mutant, coincided with the appearance of additional oligosaccharide bands not observed in WT. Asterisks (*) indicate these novel bands.

To investigate whether AtXYXT1 also adds xylosyl side chains two residues from GlcA substitutions in xylan, we analysed basal stem AIR from *esk1 kak8* plants ectopically expressing AtXYXT1 under the *AtIRX3* promoter. Samples were digested with GH30 followed by GH3 xylosidase, and compared to the *esk1 kak8* control. As observed with PmXYXT2, GH30 digestion of pIRX3::AtXYXT1 lines produced several additional bands appearing between the UXₙ ladder. Subsequent GH3 treatment caused these bands to collapse predominantly into a single, GH62-sensitive oligosaccharide migrating in the Y^2^XXXUX region (Figure 5B). To confirm its identity, the oligosaccharide was 2-AA labelled, and analysed by MALDI-CID mass spectrometry (Figure 5C). The resulting fragmentation pattern confirmed a structure containing one GlcA and a single pentose substitution at the reducing end of the molecule. GH3 digestion yielded an oligosaccharide cleaved up to the terminal xylose carrying the pentose, consistent with a β-1,2-linkage at the reducing end. Together, these results confirm the presence of a β-1,2-linked xylosyl side chain and support the formation of a regularly substituted structure such as Y^2^XXXUX. This demonstrates that AtXYXT1 generates the same GlcA-linked xyolsylated xylan motif observed in Arabidopsis primary walls when expressed in SCW tissue. Notably, AtXYXT1 activity was markedly reduced when expressed in SCW WT plants, similar to PmXYXT2, suggesting that secondary wall acetylation hinders its activity (Supporting Figure S5B).

### Arabidopsis XYXT triple mutants lack xylosylated xylan and show tissue specific developmental defects

To evaluate the contribution of endogenous AtXYXT1 to xylan β-1,2-xylosylation, we analysed AIR from Arabidopsis callus of an *atxyxt1* T-DNA knockout line. GH30 and GH3 digestion revealed that the Y^2^XXXUX oligosaccharide was still detectable in *atxyxt1*, although slightly reduced compared to WT. As AtXYXT1 seems not the only enzyme responsible for the synthesis of Y^2^XXXUX, we tested for redundancy with the two closest homologues, named AtXYXT2 and AtXYXT3 (Figure 2B). We identified a knock-out T-DNA insertion mutant for *AtXYXT2* (Supporting Figure S5) and generated the *atxyxt1 atxyxt2* double mutant. In this background, Y^2^XXXUX still remains detectable, suggesting involvement of AtXYXT3. As T-DNA homozygous knockout lines for *AtXYXT3* could not be obtained, we used CRISPR-Cas9 to target AtXYXT3 in the *atxyxt1 atxyxt2* double mutant background. In the resulting triple mutant *atxyxt1 atxyxt2 atxyxt3*, the Y^2^XXXUX oligosaccharide was absent (Figure 5D), providing genetic evidence that these three genes act redundantly to generate the β-1,2-xylosylated structure in Arabidopsis primary wall xylan.

Notably, reduction of Y^2^XXXUX in the single and double mutants, and its complete loss in the triple mutant, coincided with the appearance of additional oligosaccharide bands not observed in WT (*) (Figure 5D). These likely reflect additional xylan substitutions that are typically masked by 2-xylosylation, hinting at additional structural variation in primary wall xylan that remains to be discovered.

AtXYXT1 (MUCI21/MUM5) has previously been implicated in seed coat mucilage adherence, with *atxyxt1* (*muci21/mum5*) mutants showing a pronounced reduction in mucilage adherence (Voiniciuc *et al*., 2015; Ralet *et al*., 2016). To investigate whether other AtXYXT genes contribute to this phenotype, we examined the mucilage of *atxyxt2*, the *atxyxt1 atxyxt2* double mutant, and the *atxyxt1 atxyxt2 atxyxt3* triple mutant. While *atxyxt2* seeds mucilage appeared similar to WT, both the double and triple mutants showed mucilage phenotypes comparable to *atxyxt1* alone (Supporting Figure S7B). These findings indicate that AtXYXT1 accounts for most of the effect on mucilage adherence, in line with its predominant expression in the seed coat (Supporting Figure 7A).

To investigate whether *AtXYXT* genes might also play roles later in development, we examined leaf senescence, where transcriptome data show elevated expression of all three *AtXYXT* genes (Supporting Figure S8A). When grown under standard long-day conditions, *atxyxt* mutants displayed visible differences in rosette appearance eight weeks after germination. While WT plants exhibited clear signs of senescence—such as widespread leaf yellowing—*atxyxt1* and *atxyxt1 atxyxt2* mutants remained greener and overall healthier, with the triple mutant *atxyxt1 atxyxt2 atxyxt3* showing the most pronounced delay in senescence (Supporting Figure S8B). Although these differences were not evident at earlier timepoints, they coincide with the temporal expression pattern of *AtXYXT* genes (Supporting Figure S8A), suggesting a potential role for β-1,2-xylosylated xylan in cell wall remodelling during ageing.

## Discussion

In this study, we identify a previously uncharacterised β-1,2-linked xylosyl side chain on xylan, forming a distinctive XUXY^2^XX oligosaccharide after xylanase GH11 hydrolysis. This structure was consistently detected in *Metasequoia*, *Pinus*, *Pseudotsuga*, and *Arabidopsis*, and in each case was enriched in primary wall-rich tissues such as needles and PEM. Its detection across both gymnosperms and angiosperms suggests that this xylosyl side chains on xylan are evolutionarily conserved and predates the divergence of seed plant lineages. Notably, this structure was markedly reduced or undetectable in mature, lignified tissues, supporting its specific association with primary cell walls.

Structural characterisation of XUXY²XX was enabled by an unexpected enzymatic signature: the oligosaccharide was resistant to GH51 arabinofuranosidase but selectively cleaved by *Pa*GH62, a GH62 enzyme typically associated with arabinofuranose removal. MALDI-CID mass spectrometry and NMR confirmed that the side chain is a β-1,2-linked xylose, not arabinose— revealing β-1,2-xylosidase activity under our assay conditions. To the best of our knowledge, such activity has not been previously reported in GH62 enzymes. This finding enabled the identification of a novel xylan side chain.

The evolutionary origins and diversity of pentosyl side chains on xylan have remained unclear, but growing evidence suggests that these modifications are both ancient and structurally varied across plant lineages. In *Physcomitrium*, mass spectrometry has revealed pentosyl substitutions directly on the xylan backbone, suggesting that such modifications were already present in early land plants (Kulkarni *et al*., 2012). While the precise identity of the pentose remains unresolved, arabinosyl side chains are common across plant species—particularly 3-linked arabinoses in commelinid monocots—and 2-linked arabinosyl residues have also been reported (Naran *et al*., 2008; McCleary *et al*., 2015; McCleary *et al*., 2015).

By contrast, evidence for xylosyl side chains is sparse and mostly limited to early studies on *Plantago ovata* husk xylan, where 2- or 3-linked xylose substitution have been proposed (Samuelsen *et al*., 1999; Fischer *et al*., 2004; Edwards *et al*., 2003; Guo *et al*., 2008; Yin *et al*., 2012). In *Arabidopsis*, linkage analysis has similarly suggested 2-linked residues in seed mucilage (Voiniciuc *et al*., 2015; Ralet *et al*., 2016), which may indicate xylosyl substitution. However, linkage analysis alone cannot definitively confirm the identity of the substituent sugar or its position within the polymer. The first β-1,2-xylosyltransferase activity was characterised in rice (OsXYXT1; Zhong *et al*., 2018), though the corresponding structure *in planta* has not been identified in grass xylan. In conifers, two *Pinus taeda* xylan xylosyltransferases (PtXYXT1 and PtXYXT2) also displayed β-1,2-xylosyltransferase activity *in vitro*, and a minor NMR signal at 4.64 ppm, attributed to H1 of 2-O-xylosylated residues, hinted at the presence of such modifications in pine wood xylan (Zhong *et al*., 2022). However, no structural analysis confirmed the precise position of these branches nor their relationship to GlcA substitutions.

Our identification of the XUXY^2^XX oligosaccharide in both conifer and Arabidopsis primary cell walls suggests that β-1,2-xylosylation of xylan may not be more common than previously reported. This modification appears specific to primary walls, where xylan is less abundant and structurally harder to resolve, potentially explaining why it has remained undetected. Although β-1,2-xylosyltransferase activity has been demonstrated for two conifer GT61 groups, our results reveal a key difference in substrate specificity: group III enzymes consistently position the xylosyl branch two residues upstream of GlcA. We find the same positional precision in Arabidopsis AtXYXT1, and genetic evidence confirms that the *atxyxt1 atxyxt2 atxyxt3* triple mutant lacks this modification entirely. These findings establish β-1,2-xylosylation as a conserved and positionally specific structure of primary cell wall xylans in both conifers and potentially angiosperms.

Although β-1,2-xylosylation appears to be a conserved modification of primary cell wall xylans in both conifers and angiosperms, it is not a widespread or highly abundant modification. Instead, its presence is restricted to specific tissues and developmental stages. This is consistent with the expression patterns of *AtXYXT* genes, which are not broadly expressed but instead show sharp spatial and temporal specificity. For example, *AtXYXT2* is strongly induced during heat stress and has been identified as a heat priming-associated gene (Alshareef *et al*., 2022), suggesting that β-1,2-xylosylation may play a role in stress-induced wall remodelling. Similarly, the delayed senescence and altered rosette growth observed in the *atxyxt1 atxyxt2 atxyxt3* triple mutant point to a role in modulating wall properties during late development or ageing.

What may be the function of β-1,2-xylosylation in the primary wall? A clue comes from the observed role of AtXYXT1/MUCI21/MUM5 in seed coat mucilage. In *atxyxt1/muci21/mum5* mutants, pectin is secreted but fails to remain attached to the seed surface, coinciding with the loss of β-1,2-xylosylation in mucilage xylan (Voiniciuc *et al*., 2015). An alternative model proposed that AtXYXT1/MUCI21/MUM5 modifies RG-I to generate xylosylated side chains involved in xylan-RGI adsorption (Ralet *et al*., 2016; Saez-Aguayo and Largo-Gosens, 2022), but our data demonstrate direct modification of xylan. We detect a single β-1,2-linked xylose branch on the xylan backbone, consistent with AtXYXT1 acting as a xylan β-1,2-xylosyltransferase, suggesting that the xylose side chain on xylan plays a role in preventing detachment of pectin in the seed mucilage.

This interpretation is consistent with the presence of covalently linked xylan–pectin complexes in Arabidopsis and conifers. In Arabidopsis, the APAP1 proteoglycan links glucuronoxylan and pectin into a tri-polymeric complex in primary walls (Tan *et al*., 2013; Tan *et al*., 2024), and similar assemblies have been described in coniferous tissues (Shakhmatov and Makarova, 2025). Notably, the XUXY^2^XX structure we identify is evenly substituted, meaning it has side chains spaced every two residues. This patterning is known to enable xylan to adopt a twofold screw conformation and bind to cellulose microfibrils (Grantham *et al*., 2017), suggesting that β-1,2-xylosylated xylan may help tether xylan–pectin assemblies to cellulose.

One possibility is that this xylosylation shields the xylan backbone from cleavage by endogenous endoxylanases. In its absence, xylan–pectin linkages may be more prone to breakdown, potentially facilitating local wall remodelling. This hypothesis could also explain why mucilage appears detached in *atxyxt1/muci21/mum5* mutants lacking 2-linked xylosylation. The observation that all three *AtXYXT* genes are expressed during the later stages of leaf development suggests that 2-xylosylation may help suppress further remodelling during senescence. Consistent with this, *atxyxt1 atxyxt2 atxyxt3* mutants display delayed leaf senescence and larger rosettes, possibly reflecting a reduced capacity to limit wall remodelling during ageing.

Although the discovery of the β-1,2-xylosylated XUXY^2^XX marks a significant advance in our understanding of primary cell wall xylan, it may only reveal part of a more complex picture. In *atxyxt* single, double, and triple mutants, the reduction and loss of β-1,2-xylosylation allows additional oligosaccharides to become visible in PACE analysis that are not detected in WT (Figure 5). These previously masked structures suggest that Arabidopsis primary wall xylan may possess further, as-yet uncharacterised modifications, and point to structural complexity beyond the conserved GlcA–Xyl pattern described here.

These findings define a conserved structural feature of PCW xylan and point to a broader, still emerging picture of its potential roles in wall architecture. β-1,2-xylosylation may contribute to aspects of wall biology that remain poorly understood, including possible interactions with pectins, matrix assembly, or remodelling during development and ageing. Further work will be needed to explore these possibilities.

## Materials and Methods

### Plant material and growth conditions

Arabidopsis and *Nicotiana benthamiana* plants were grown in controlled conditions at 21°C under 16h light cycle on Levington M3 vermiculite soil. Liquid callus cultures were generated and maintained as described (Prime *et al*., 2000). Mutants used were in Col-0 ecotype: *gux1 gux 2* (Mortimer *et al*., 2010), *esk1 kak8* (Bensussan *et al*., 2015), *muci21-1* (At3g10320, Salk_ 041744;(Voiniciuc *et al*., 2015)). *atxyxt2* was ordered from NASC (At2g41640, Salk_108485; (Scholl *et al*., 2000), Col-0 was used as wildtype control (WT). Homozygosity of the T-DNA knockout line for *atxyxt2* was confirmed by RT–PCR as described (Temple *et al*., 2019). Allelic triple mutants were generated using CRISPR–Cas9 constructs targeting *atxyxt3* in the *atxyxt1 atxyxt2* double mutant background, and mutations were verified by Sanger sequencing (for oligonucleotide sequences, see Supplementary Table 1).

Gymnosperm tissue samples were primarily obtained from *Pinus radiata* (D. Don). *P. radiata* tissue samples (vegetative bud, phloem, needle, root, herbaceous stem and cambium/xylem) were collected from four individual 6-year-old clonal trees grown outdoors in Rotorua, New Zealand. PEM cultures were established from immature embryos excised from green cones and maintained on Glitz2 medium (Hargreaves et al., 2017). All samples were snap-frozen in liquid nitrogen and stored at −80 °C for subsequent mRNA extraction and AIR preparation. Additional material from *Metasequoia glyptostroboides*, *Picea abies*, and *Pseudotsuga menziesii* was also included for comparative analysis.

### Molecular cloning and plant transformation

Coding sequences (CDSs) of *Pseudotsuga menziesii* PmXYXT1 and PmXYXT2, *Oryza sativa* OsXYXT1, and *Arabidopsis thaliana* AtXYXT1 (At3g10320) were synthesised (Genewiz) and domesticated for Golden Gate assembly. Domestication involved removing internal BsaI and BpiI recognition sites—used as type IIS restriction enzymes in the OpenPlant common syntax system—via synonymous codon substitutions without altering the amino acid sequence. Each CDS was assembled under the *IRX3* promoter into binary vectors using Golden Gate cloning (Patron *et al*., 2015). Constructs were introduced into *A. thaliana* Col-0, *gux1 gux2*, and *esk1 kak8* backgrounds by the floral dip method (Clough and Bent, 1998). T_3_ homozygous transgenic lines were selected using the Fast Green screenable marker system (Shimada *et al*., 2010).

For CRISPR experiments, a single sgRNA targeting *AtXYXT3* was designed using CHOPCHOP (https://chopchop.cbu.uib.no) and cloned into a binary vector following the Golden Gate MoClo Toolkit protocol (Lawrenson *et al*., 2015).

Domesticated CDSs fused to a C-terminal myc tag were cloned into NruI-digested pEAQ-HT vectors (Sainsbury *et al*., 2009) using T4 DNA ligase. Additionally, PmXYXT2-GFP fusions driven by the constitutive 35S promoter were assembled via MoClo and used for subcellular localisation studies in *Nicotiana benthamiana*.

### Enzyme digestions

In most experiments, hemicelluloses were extracted from AIR prepared from Arabidopsis basal stem, leaf, callus, and conifer tissues by incubation with 4 M NaOH at room temperature for 1 hour. After centrifugation, soluble fractions were desalted and buffer-exchanged using PD-10 columns equilibrated with 50 mM ammonium acetate (pH 6.0). Aliquots (typically 250 µL) were subjected to enzymatic digestion and analysis.

NpXyn11A xylanase (GH11) (Megazyme; 30 U per reaction) was used for overnight digestion at 30 °C to hydrolyse accessible xylan. Where applicable, additional enzymes were applied as follows: *Bacteroides ovatus* GH115 glucuronidase (3 µL of 1 mg/mL; NZYtech), *Meripilus giganteus* GH51 α-arabinofuranosidase (3 µL of 3.5 mg/mL), and *Penicillium aurantiogriseum* GH62 α-arabinofuranosidase (3 µL of 3.5 mg/mL). The GH51 and GH62 enzymes were kindly provided by Dr Kristian Krogh (Novozymes).

In a separate experimental series, *Erwinia chrysanthemi* GH30 glucuronoxylanase (0.5 µL of 16 mg/mL) was incubated for 1 hour at room temperature. Where applicable, GH3 β-xylosidase was added following GH30 digestion. Prior to GH62 treatment, the GH30 and GH3 enzymes were removed by filtration using 10 kDa Nanosep centrifugal filters.

### PACE analysis

PACE was performed as described (Yu *et al*., 2024). Dry samples and xylo-oligosaccharide standards were fluorescently labelled by reductive amination using 8-aminonaphthalene-1,3,6-trisulfonic acid (ANTS) and 2-picoline borane. A labelling mix was prepared by combining 0.2 M ANTS (in 3:17 acetic acid:H₂O), 0.2 M 2-picoline borane (in DMSO), and derivatisation buffer (H₂O:DMSO:acetic acid, 17:20:3, v/v) in equal volumes. Twenty microlitres of labelling mix were added to each dry sample, and reactions were incubated at 37 °C overnight. Samples were then dried in a vacuum concentrator at 60 °C and resuspended in 30 µl of 3 M urea.

Labelled oligosaccharides were resolved by polyacrylamide gel electrophoresis in 0.1 M Tris-borate buffer (pH 8.2) at 1,000 V for 1 hour at 10 °C. Gels were imaged using a G-Box CCD system under 365 nm UV illumination.

### GT61 gene identification and phylogenetic analysis

Putative *P. radiata* GT61 transcripts were collected from in-house *P. radiata* transcriptomes via similarity searches against known *A. thaliana* GT61 genes. Consensus coding sequences were then generated via multiple sequence alignment (Clustal Omega) and mapped to the *P. radiata* genome (Sturrock *et al*., 2025) using the minimap2 function in Geneious (Siemens Digital Industries Software) to retrieve their genomic context and further define consensus sequences. Full-length genes were recovered from *P. radiata* xylem mRNA using PCR and cloned into plasmids for sequence verification.

For the GT61 phylogenetic analysis, amino acid sequences from *Pseudotsuga menziesii*, *Picea abies*, and *Pinus taeda* were obtained from the PLAZA Gymno01 database (Proost *et al*., 2015); https://bioinformatics.psb.ugent.be/plaza/versions/gymno-plaza/) and included the newly identified *P. radiata* GT61 sequences. Sequences from *A. thaliana*, *Oryza sativa*, *Physcomitrella patens*, *Marchantia polymorpha*, and *Brachypodium distachyon* were retrieved from PLAZA 5.0 (Van Bel *et al*., 2022). Only full-length protein sequences were included in the analysis. Sequences were aligned using MUSCLE (Multiple Sequence Comparison by Log-Expectation), and phylogenetic reconstruction was performed using the maximum likelihood method with 100 bootstrap replicates in MEGA version 10. The resulting tree was visualised using FigTree version 1.4.2.

### *Pinus radiata* RNA isolation and cDNA synthesis

RNA extractions were performed on 50 mg of plant material from each tissue type using protocol B of the Spectrum Plant Total RNA kit (Sigma Aldrich, St. Louis, MO, USA). Concentration, purity and integrity of total RNA were determined fluorometrically (Qubit, Thermo Fisher Scientific, Waltham, Massachusetts, USA). Reverse transcription (RT) was carried out using the iScript gDNA clear cDNA synthesis kit (Bio-Rad, Hercules, CA, USA) with 900 ng of DNase-treated total RNA as a template.

### Droplet digital PCR (ddPCR) analysis of *PrGT61* family expression

The ddPCR was performed using the Bio-Rad QX200 system (Bio-Rad, Hercules, CA, USA). Each reaction contained 11 μL of ddPCR EvaGreen Supermix (Bio-Rad), primers at a final concentration of 100 nM each, 4 μL of a 1:20 cDNA template dilution and ultrapure water to a final volume of 22 μL. Droplet generation was carried out according to manufacturer’s instructions (Bio-Rad) and droplets were quantified in a QX200 Droplet Reader (Bio-Rad).

Results were analysed with QX Manager Software 2.3 Standard Edition. No amplification was observed from RNA-only controls (no RT).

### Microsome preparation and *in vitro* assay

Approximately 15 g of *Nicotiana benthamiana* leaves expressing GT61 constructs in the pEAQ-HT *Nicotiana benthamiana* overexpression vector were homogenised in 25 mL of ice-cold extraction buffer (50 mM HEPES-KOH pH 7.5, 0.4 M sucrose, 10 mM MgCl₂, 1 mM EDTA, 5 mM DTT, protease inhibitors). The homogenate was filtered and centrifuged at 3,500 g and 10,000 g (10 min each, 4 °C), and microsomes were pelleted at 100,000 g (1 h, 4 °C). The pellet was resuspended in 300 µL of extraction buffer, homogenised, aliquoted, and stored at –80 °C.

For in vitro assays, dried AIR or acetylated xylan was resuspended in 30 µL of master mix (0.5 mM DTT, 10 mM MnCl₂, 10 mM MgCl₂, 2% Triton X-100, 2 mM UDP-xylose), and mixed with 30 µL of microsomes (∼500 µg protein). Reactions were incubated overnight at room temperature, then terminated by heating at 100 °C for 10 min. Polysaccharides were extracted by chloroform–methanol partitioning, precipitated with ethanol (–20 °C, 90 min), washed twice in 100% ethanol, and dried under vacuum (∼90 min, 45 °C).

### Size exclusion chromatography of xylan oligosaccharides from Metasquoia needles

Hemicelluloses were extracted from 500 mg from AIR prepared from Metasequoia needle with 4 M potassium hydroxide at room temperature for 1 hour. After centrifugation, the extracted hemicelluloses in the supernatant were passed through a PD-10 column (Cytiva) for solvent exchange to 50 mM ammonium acetate buffer pH 6.0. The xylan in the sample was then hydrolysed by an GH11 (600 U, Megazyme) and MgGH51 α-arabinofuranosidase (216 µg, Novozyme) at 37°C overnight. To reduce the viscosity, the undigested polysaccharides were precipitated by ethanol and xylan oligosaccharides in the supernatant were lyophilised. The dried oligosaccharides were dissolved into 5 mL of the buffer and heated at 80°C for 10 min. The sample was then applied to a Bio-Gel P-2 column (534 ml, φ2.5×104 cm) equilibrated with the buffer, and then eluted by gravity with a flow rate of 0.13 ml/min. The eluent was fractionated, and 50 µL of the fractions (2.5 ml) were subjected to PACE to monitor the elution of the xylan oligosaccharides. The purified oligosaccharides were pooled, lyophilised and stored at ─20°C.

### Oligosaccharide purification and mass spectrometry

Following enzymatic digestion, residual peptides and proteins were removed using Sep-Pak C18 reverse-phase cartridges (Thermo Scientific), as previously described. Purified oligosaccharides were labelled by reductive amination with either 2-aminobenzoic acid (2-AA) or procainamide hydrochloride (ProA) and subsequently desalted using GlycoClean S cartridges (ProZyme), as described. Labelled samples (1 μl) were spotted onto a MALDI plate together with 1 μl of 2,5-dihydroxybenzoic acid (DHB) matrix (10 mg ml⁻¹ in 50% aqueous methanol) and analysed by MALDI-time-of-flight/time-of-flight mass spectrometry (ToF/ToF– MS) using an UltrafleXtreme (Bruker), as described (Feijao *et al*., 2022).

### Solution NMR analysis

The Metasequoia xylan oligosaccharide enriched by size exclusion chromatography was solubilized in 600μL of D_2_O and analysed at 298K on an 800MHz Bruker AVANCE III spectrometer fitted with a TXI CryoProbe. In addition to one dimensional *1*H spectra, both homonuclear – *1*H,*1*H-COSY, *1*H,*1*H-TOCSY, and *1*H,*1*H-NOESY – and heteronuclear – *1*H,*13*C-HSQC, *1*H,*13*C-HSQC-TOCSY, and *1*H,*13*C-H2BC – two-dimensional spectra were recorded. Mixing times of 120 milliseconds, 200 milliseconds, and 100 milliseconds were used for the 2D *1*H,*1*H-TOCSY, *1*H,*1*H-NOESY, and *1*H,*13*C-HSQC-TOCSY respectively. Spectra were processed using Bruker Topspin 3.1 and assigned using the Collaborative Computing Project for NMR Analysis 2.4.2 software (Vranken *et al*., 2005).

## Acknowledgements

We would like to thank Scion staff: Barbara Geddes and Cathie Reeves for the PEM creation and growth, Lorelle Phillips and Heather Flint for the samples collection, Yeganeh Eslami for the AIR preparation and Tancred Frickey for the genome mining of radiata pine.

## Author contributions

H.T., N.A., and P.D. conceptualised the research and manuscript. H.T. performed the majority of the molecular genetics and biochemical experiments, with assistance from Y.Y., K.D., H.Y., A.L., A.E.P., L.Y., X.Y., and N.A. Mass spectrometry (MS) experiments were conducted by T.T. and J.R.W., and NMR experiments were performed by J.A.L. and K.S.

M.S. contributed to project development and coordinated the *Pinus radiata* sample collection, gene identification, cloning, sequencing, and RNA-based analyses (RNA extraction, RT-PCR, ddPCR), which were carried out in collaboration with C.F., K.H., and G.T.; N.S. and K.B.K. produced the *Pa*GH62. The manuscript was written by H.T., M.S., N.A., and P.D., with input from all authors. P.D. and M.S. acquired funding.

## Data availability statement

All relevant data are included within the manuscript and its supplemental files.

## Supporting figure legends

**Supporting Figure S1:**
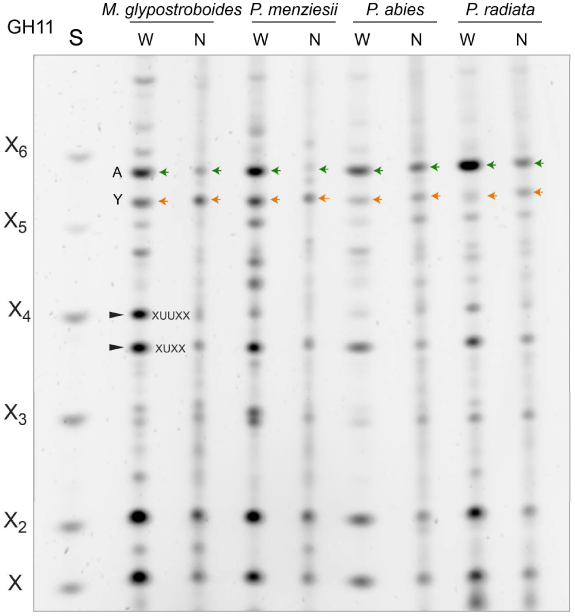
Difference in xylan structure in needles (N) and wood (W) PACE analysis of AIR from *Pinus radiata*, *Pinus abies*, *Pseudotsuga menziesii* and *Metasequoia glyptostroboides* hydrolysed with xylanase GH11. Note the difference in ratio of unknown oligosaccharide A and Y. S: Standard of xylo-oligosaccharides X_1_-X_6_.

**Supporting Figure S2:**
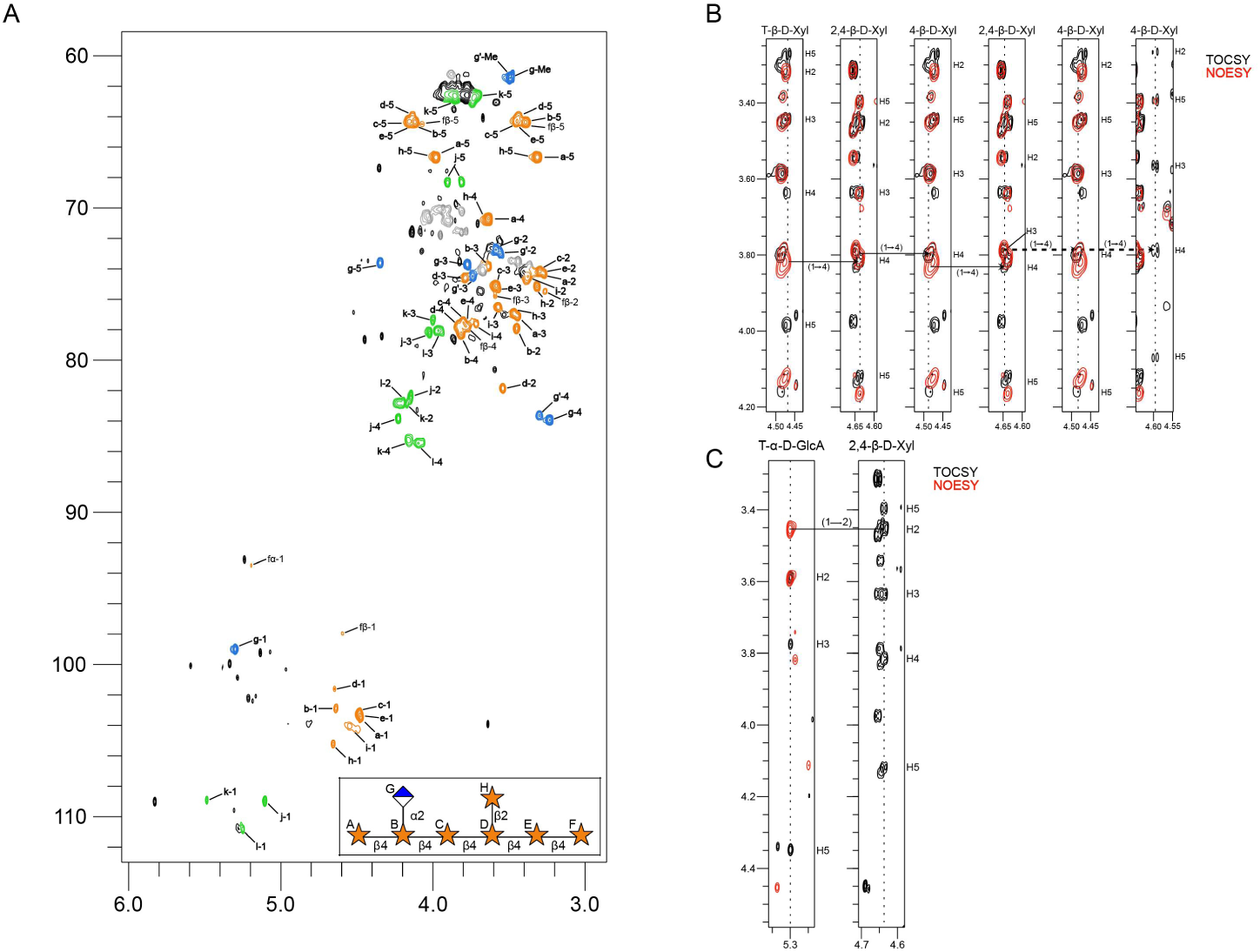
Structural analysis of Y xylo-oligosaccharide from Metasequoia needles. **A** 2D *1*H,*13*C-HSQC spectrum of a metasequoia xylan oligosaccharide obtained at 800MHz in D*_2_*O at 298 K. Letters a – h correspond to the residues labelled in the Symbol Nomneclature For Glycans (SNFG) structure shown in the insert, bottom right, while i corresponds to a further β-D-xylose residue and j – l to arabinose residues, where there was no linkage evidence for any of these additional residues to the structure identified. α and β indicate the reducing end anomer conformation that the corresponding peak relates to. **B** NMR analysis of the T-β-D-Xyl-(1 → 4)-β-D-Xyl-(1 → 4)-β-D-Xyl-(1 → 4)-β-D-Xyl-(1 → 4)-β-D-Xyl-(1 → 4)-β-D-Xyl oligosaccharide backbone structure, using *1*H strip plots for *1*H,*1*H-TOCSY (black) and *1*H,*1*H-NOESY (red) spectra to highlight the NOE connectivity arising from these glycosidic linkages. Full arrows indicate definite NOE connections, while dashed arrows indicate tentatively identified ones. **C** NMR analysis of the T-α-D-GlcA-(1 → 2)-β-D-Xyl side chain to backbone linkage, using *1*H strip plots for *1*H,*1*H-TOCSY (black) and *1*H,*1*H-NOESY (red) spectra to highlight the NOE connectivity arising from this glycosidic linkage.

**Supporting Figure S3:**
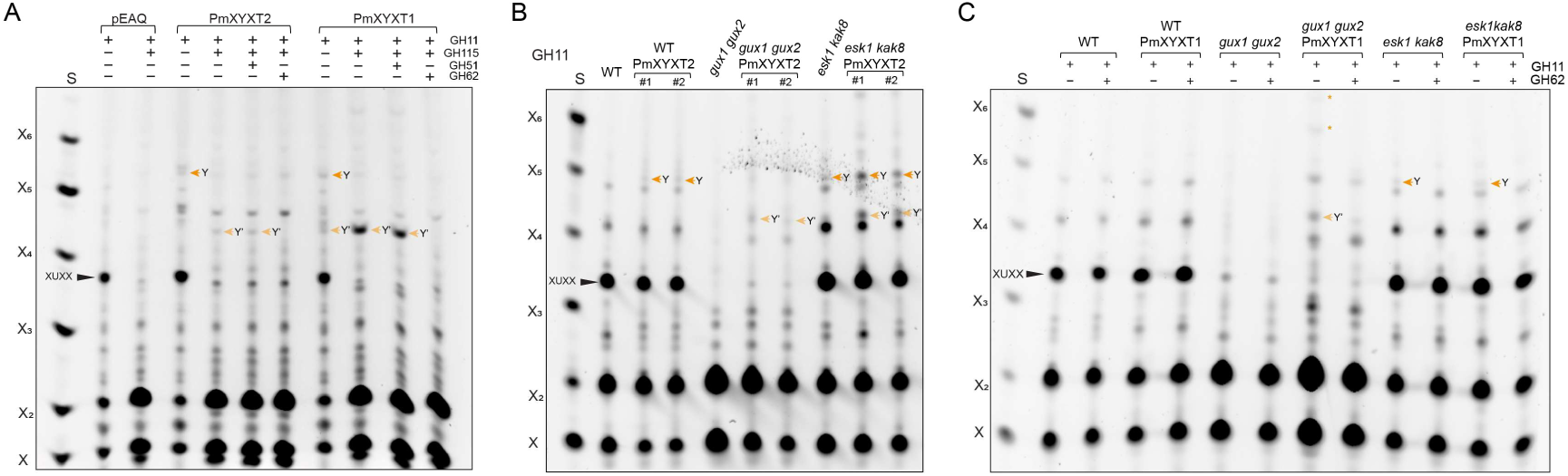
Differential activity of GT61 xylosyltransferases from *Pseudotsuga menziesii* group II (PmXYXT2) and III (PmXYXT1) **A** PACE analysis of the *in vitro* activity of PmXYXT2 and PmXYXT1 on Arabidopsis de-acetylated xylan acceptor, which was hydrolysed after the enzymatic reaction, with xylanase GH11, glucuronidase GH115, GH51 and GH62 as indicated. pEAQ: empty vector control. S: Standard of xylo-oligosaccharides X_1_-X_6_. **B** PACE analysis of xylan oligosaccharides from stem tissue heterologously expressing *pIRX3::PmXYXT2* hydrolysed by xylanase GH11. Expression in WT, *gux1 gux2* and *esk1 kak8* was analysed compared to their respective genetic background plants. #1 and #2 shows the results of two independent transgenic lines. **C** PACE analysis of xylan oligosaccharides from stem tissue heterologously expressing *pIRX3::PmXYXT1* hydrolysed by xylanase GH11 and GH62 as indicated. Expression in WT, *gux1 gux2* and *esk1 kak8* was analysed compared to untransfected plants. Note: Gels in B and C were run with 10% acrylamide (instead of the standard 20%), causing Y and Y′ to migrate slightly faster.

**Supporting Figure S4:**
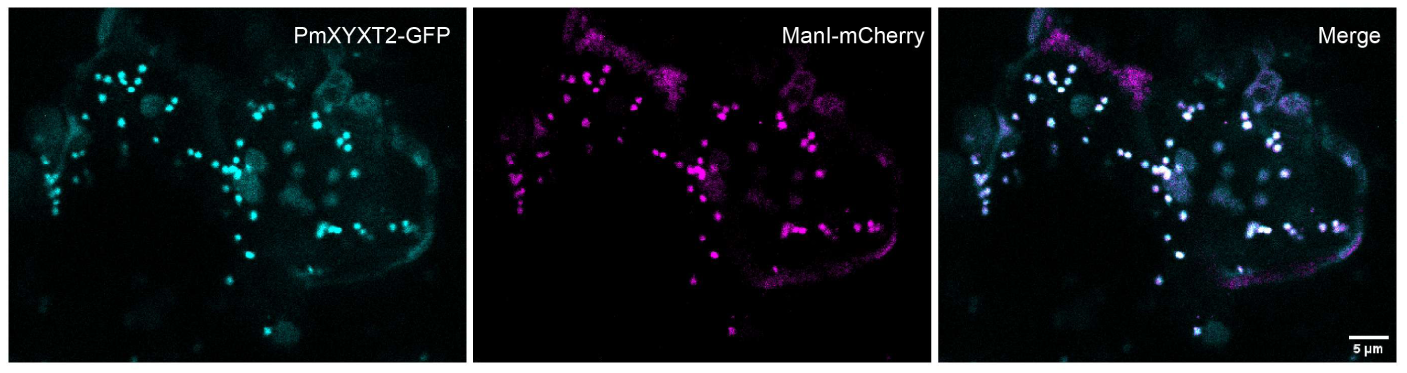
Subcellular localisation of PmXYXT2 in *Nicotiana benthamiana* leaves. PmXYXT2 localises to the Golgi apparatus. Transient co-expression of *35S::PmXYXT2-GFP* (cyan) and 35S::ManI-mCherry (*magenta*) in tobacco leaves imaged by confocal fluorescence microscopy, showing co-localisation at the Golgi in the merged image (*white*). Scale bar 5 µm.

**Supporting Figure S5:**
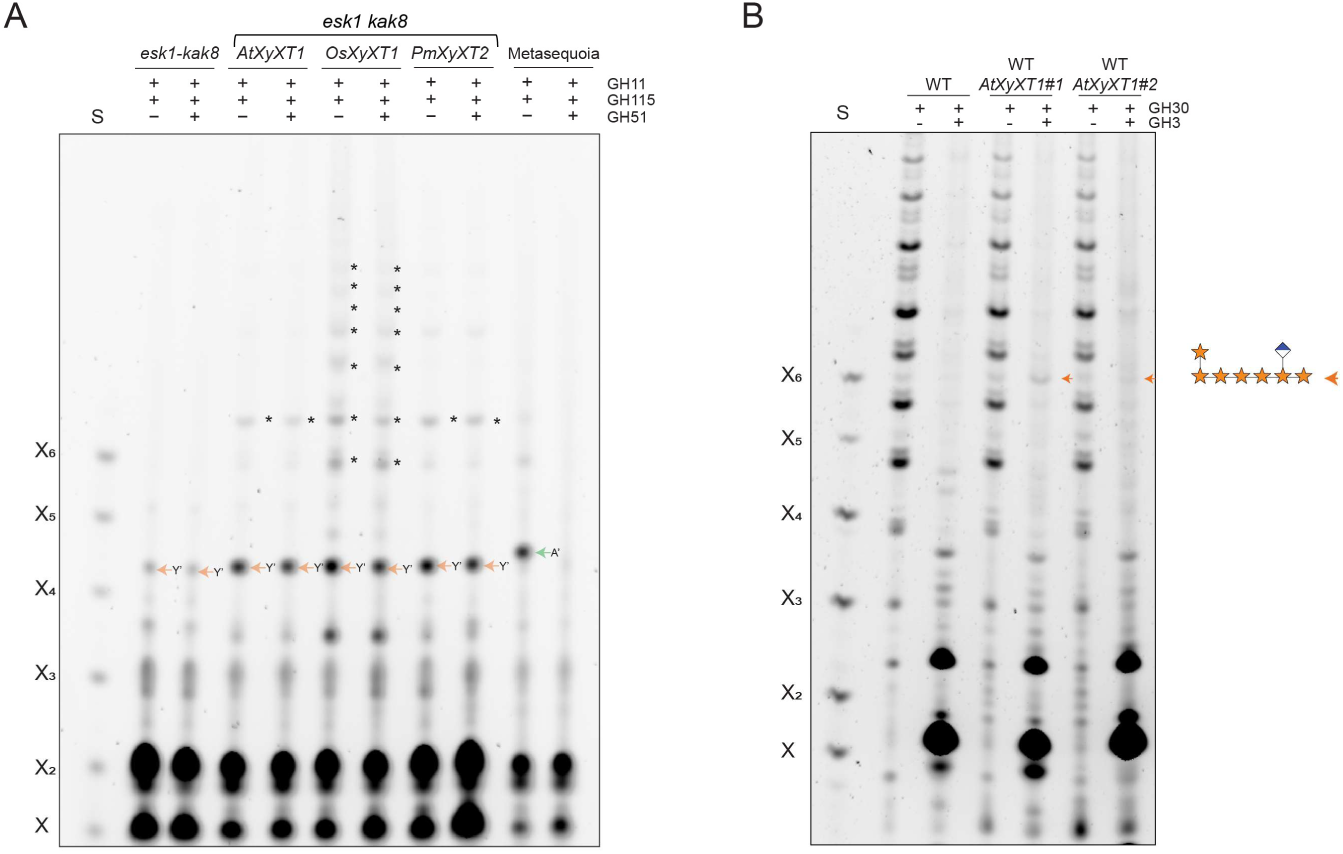
PACE analysis of xylan oligosaccharides generated by ectopic expression of GT61 xylosyltransferases in Arabidopsis stem. **A** PACE analysis of *esk1 kak8* xylan expressing AtXYXT1/MUCI21/MUM5, OsXYXT1 and PmXYXT2 under the *IRX3* promoter. AIR from stem was hydrolysed by xylanase GH11, GH115 and arabinofuranosidase GH51 as indicated. Control: *Metasequoia glyptostroboides* wood. Asterisks indicate additional, unidentified oligosaccharides that likely represent more complex xylosylated xylan structures. S: Standard is the xylose ladder X-X_6_. **B** PACE analysis of AtXYXT1 overexpression in Arabidopsis wild type hydrolysed with xylanase GH30 and GH3. YXXXUX (orange arrow).

**Supporting Figure S6:**
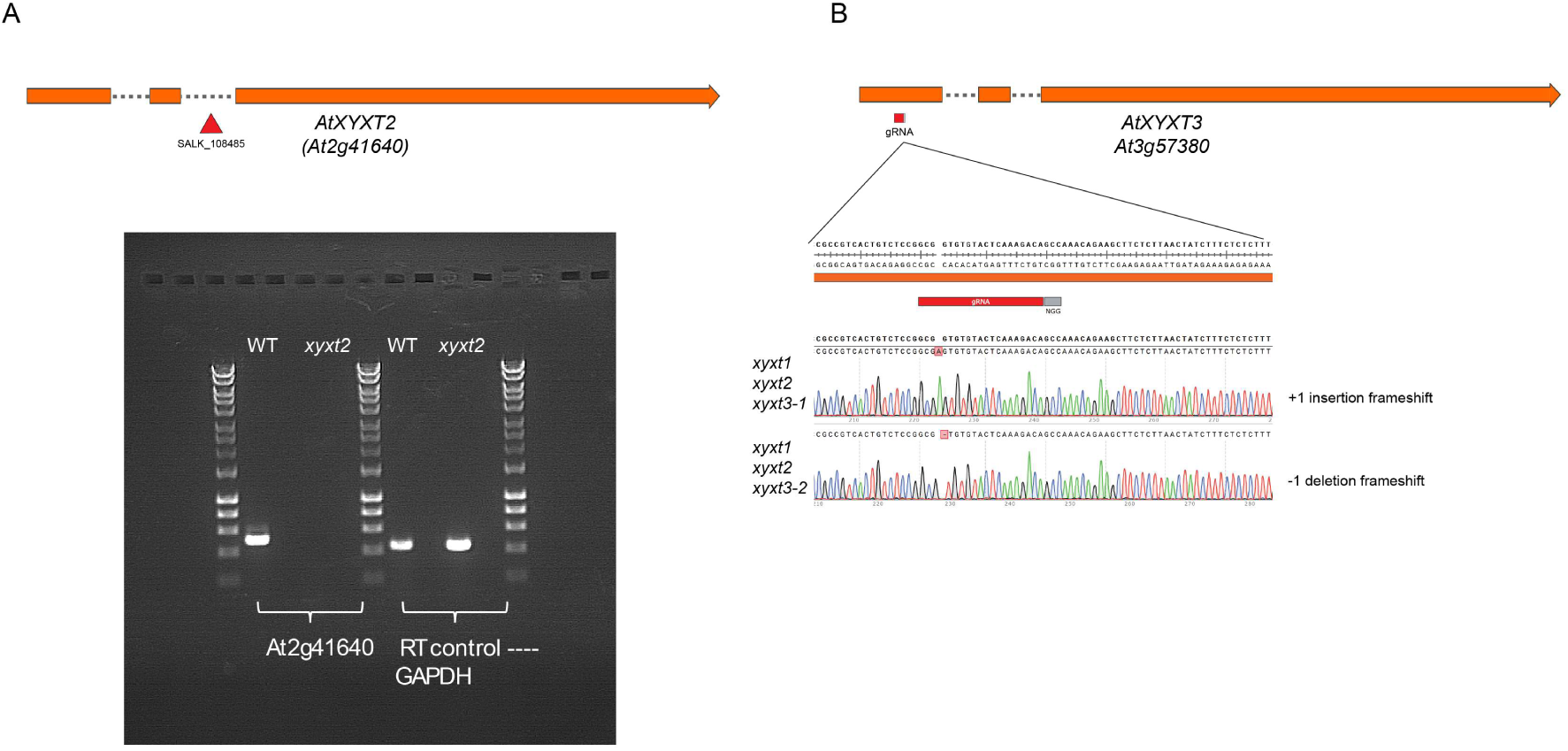
Arabidopsis AtXYXT2 and AtXYXT3 mutants. **A** Schematic representation of Arabidopsis T-DNA insertion lines of *AtXYXT2* (top) and RT-PCR confirming the knock-out via T-DNA insertion (bottom). **B** Schematic representation of CRISPR/Cas9 guide RNA (gRNA) target design (top), sequencing of CRISPR/Cas9 lines used in this analysis (bottom).

**Supporting Figure S7:**
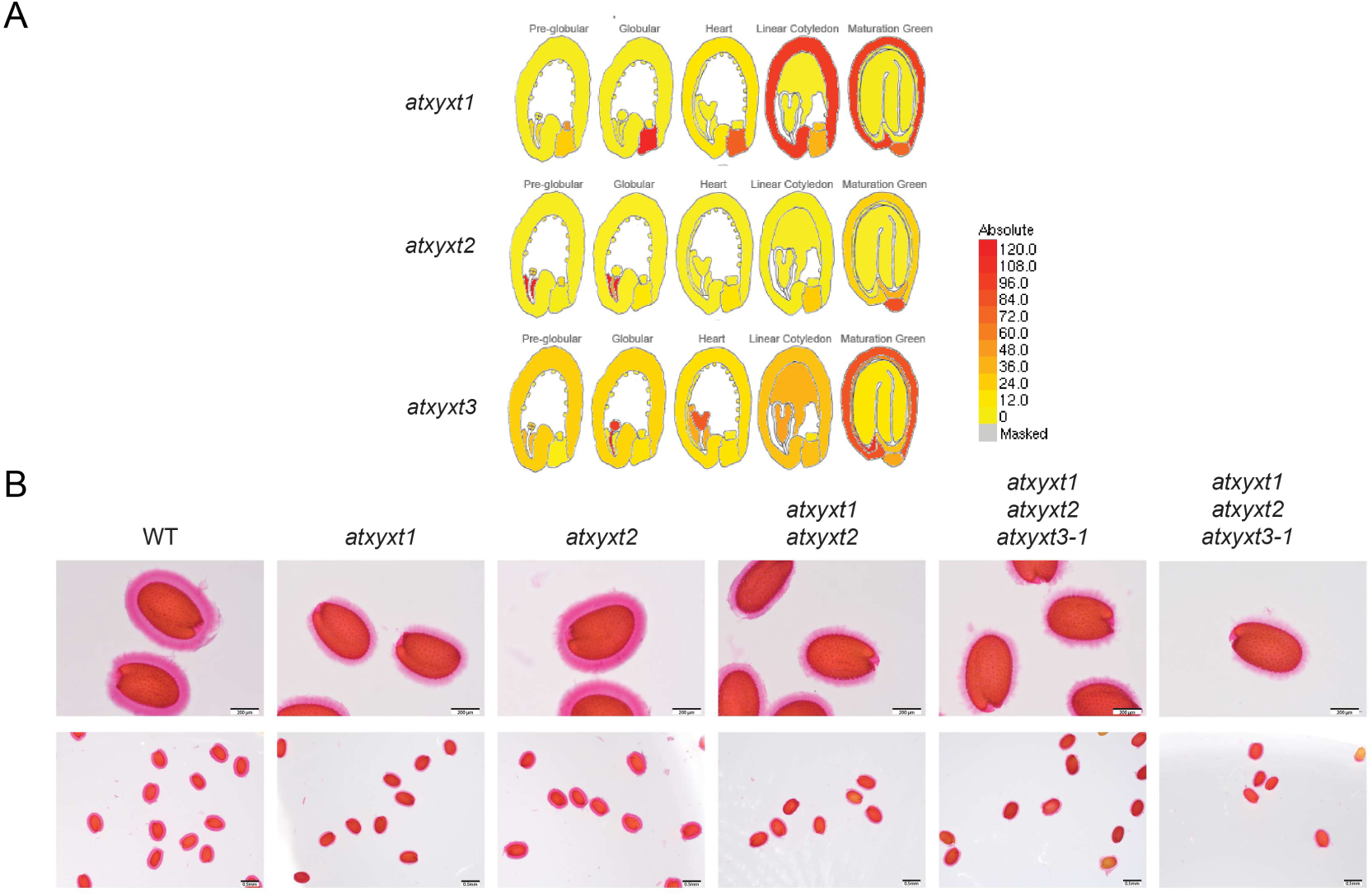
Analysis of seed phenotype in *atxyxt1 atxyxt2 atxyxt3 mutants*. **A** Seed expression of *AtXYXT1*, *AtXYXT2* and *AtXYXT3* in the seed coat obtained from eFP browser (http://bar.utoronto.ca/efp_arabidopsis/cgi-bin/efpWeb.cgi). **B** Mucilage phenotype in *atxyxt1*, *atxyxt2, atxyxt1 atxyxt2* and *atxyxt1 atxyxt2 atxyxt3*. Scale bar top panel = 200 µm, bottom panel = 500 µm. Seeds were imbibed in water, gently shaken to remove non-adherent mucilage, stained with 0.01% ruthenium red, rinsed, and imaged using a Keyence BZ-X7000 microscope.

**Supporting Figure S8:**
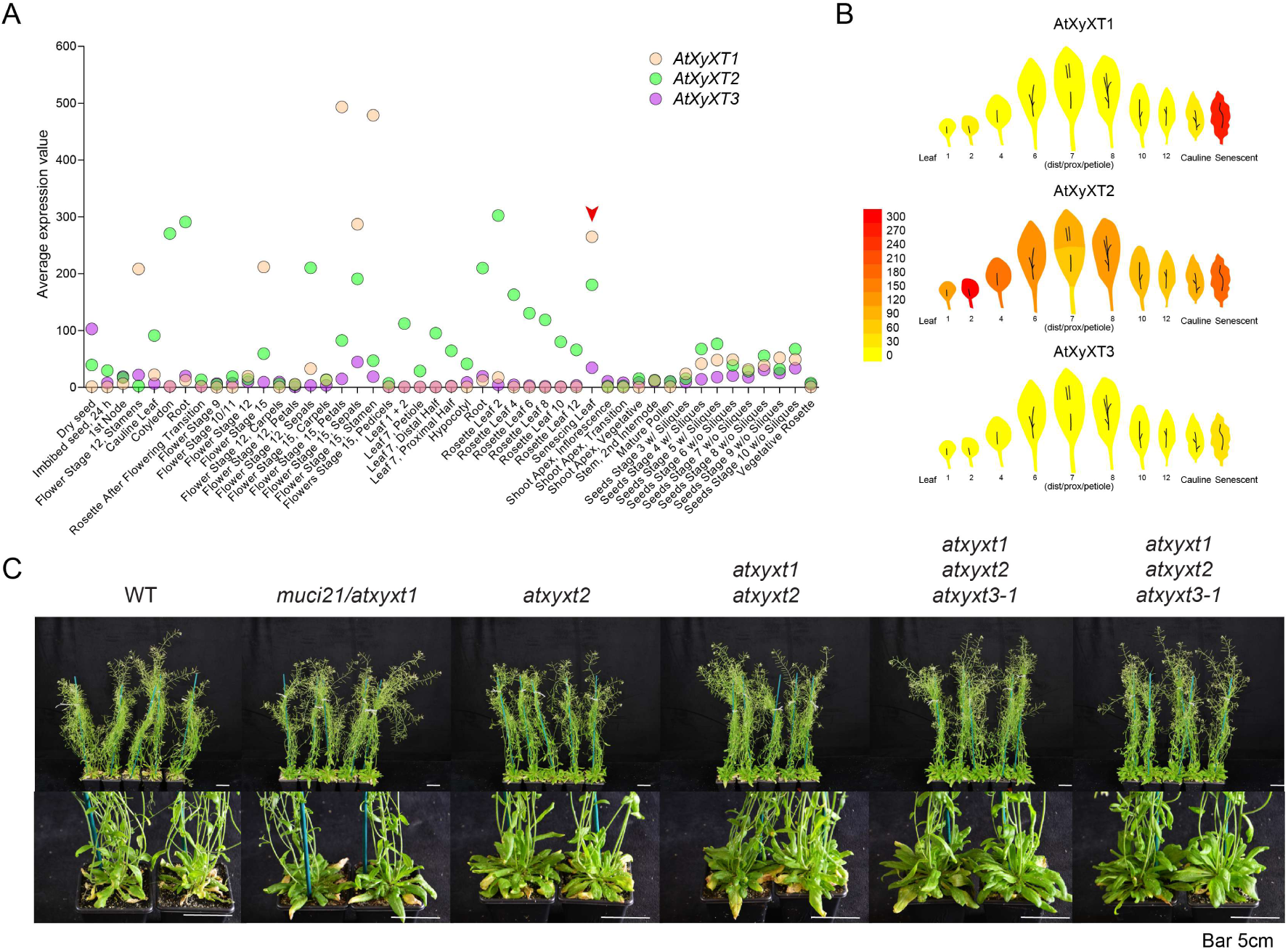
Analysis of macro-phenotype of *atxyxt1 atxyxt2 atxyxt3* triple mutants. **A** Expression data points of the *AtXYXT* family members, obtained from eFP browser. (http://bar.utoronto.ca/efp_arabidopsis/cgi-bin/efpWeb.cgi) **B** Expression heat map of *AtXYXT* family members in developing leaves, obtained from eFP browser. (http://bar.utoronto.ca/efp_arabidopsis/cgi-bin/efpWeb.cgi **C** Photographs of 8-week-old Arabidopsis plants showing wild type (WT), single mutants (*atxyxt1*, *atxyxt2*, *atxyxt3*), the double mutant (*atxyxt1 atxyxt2*), and two independent triple mutants (*atxyxt1 atxyxt2 atxyxt3-1* and *-2*). Mutant lines display visibly delayed senescence compared to WT, with leaves remaining greener for longer. The effect is subtle in single mutants, more pronounced in the double mutant, and most evident in the triple mutants, indicating a cumulative impact of *AtXYXT* gene disruption on leaf ageing and senescence timing. Scale bar = 5 cm.

<u>**Supplementary Table 1:**</u>

**Assignments Table:**

*1*H and *13*C HSQC chemical shift assignments for the metasequoia xylan oligosaccharide sample recorded. Values are given in ppm, when measured at 298 K.

**Supplementary Table 2:**
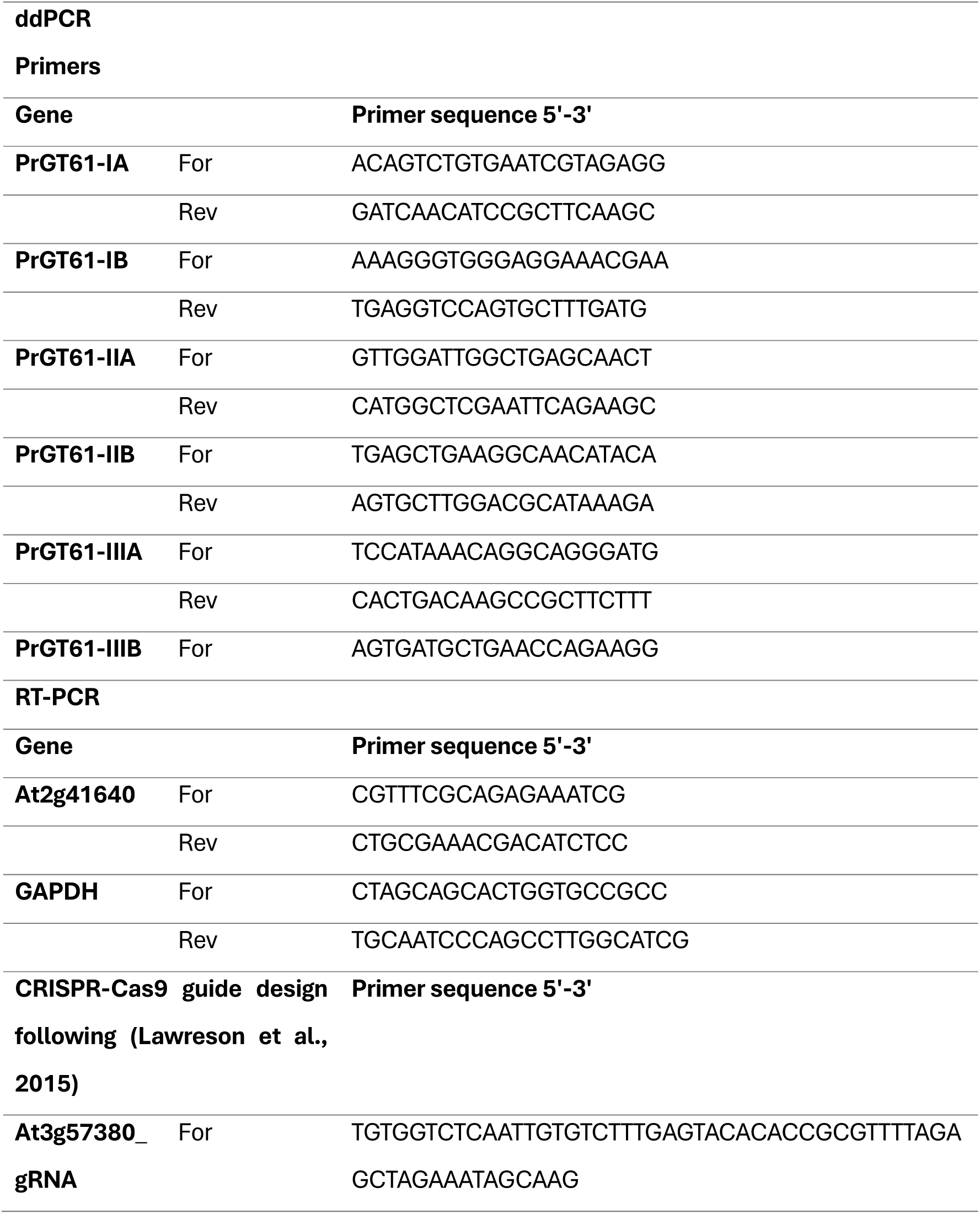
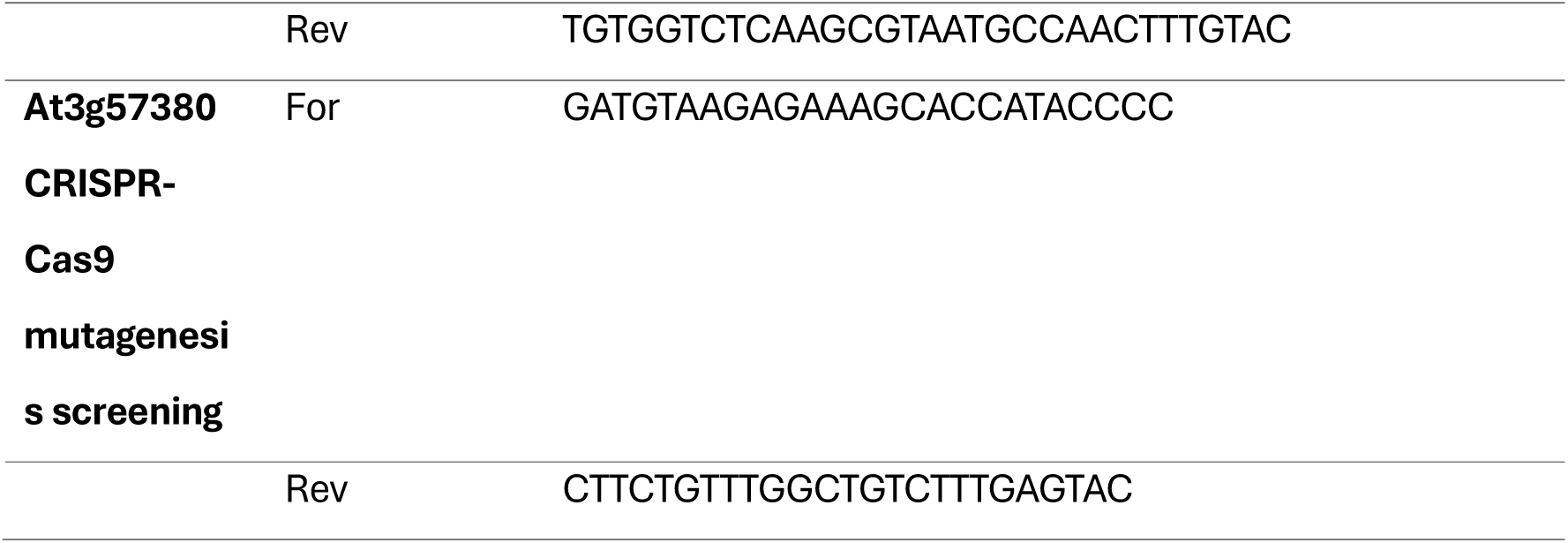
List of primers used in this study.

